# Mitochondrial heteroplasmy disrupts osteoclast differentiation and bone resorption by impairing respiratory complex I

**DOI:** 10.1101/2025.05.02.651799

**Authors:** Houfu Leng, Jiahao Jiang, Katja Gassner, Swati Midha, Raquel Justo-Méndez, Jianqing Zheng, Timothy Hall, Lin Luo, Suzanne D West, Tonia L. Vincent, Angus Wann, Kashyap A Patel, Joanna Poulton, Chris A. O’Callaghan, Ana Victoria Lechuga-Vieco, Anna Katharina Simon

## Abstract

Mitochondrial heteroplasmy, the co-existence of different mitochondrial genomes within a cell, is linked to aging and disease. Patients with heteroplasmy due to mitochondrial mutations experience multiple organ complications, particularly poor bone health and bone structure defects. However, the mechanisms involved are generally unknown, due largely to the difficulty of manipulating mtDNA in vivo. To overcome this, we leveraged a heteroplasmic mouse model and discovered that mitochondrial heteroplasmy affects a fundamental developmental process. Specifically, the differentiation of osteoclasts, which resorb bone tissue and maintain bone homeostasis. Mechanistically, there was a reduced localization of specifically respiratory complex I subunits in mitochondria in heteroplasmic mice, disrupting ATP production and osteoclast differentiation. In addition, autophagic flux is exhausted, and the autophagy inducer spermidine restores mitochondrial health and rescues osteoclast activity, both in mice and in cells from patients with primary mitochondrial disease. Together, we identify the mechanisms by which mitochondrial heteroplasmy impacts osteoclastogenesis and discover spermidine as a modulator of this process, which presents a potential treatment for human heteroplasmic conditions such as mitochondrial diseases, which are largely untreatable.

**Graphical Abstract:** 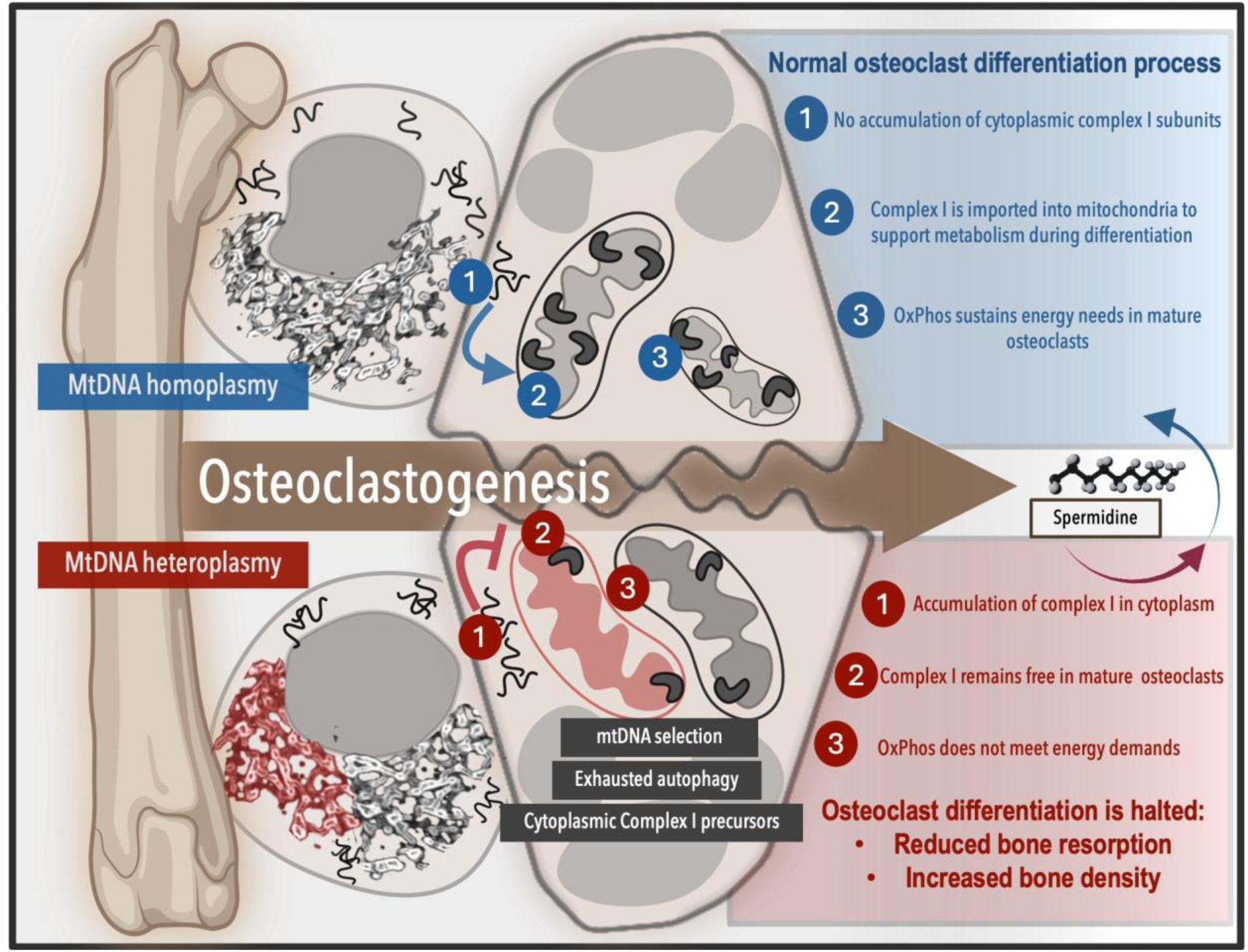

## Main

Mitochondrial heteroplasmy, the co-existence of different mtDNA variants in one cell, can lead to a wide range of pathological phenotypes, from mild effects on cellular function to severe mitochondrial diseases. MELAS syndrome (mitochondrial encephalopathy with lactic acidosis and stroke-like episodes) is a primary mitochondrial disease caused by a pathological m.3243A>G mitochondrial point mutation in the MT-TL1 gene, which was first identified three decades ago^1,2^. Primary mitochondrial diseases, including MELAS syndrome, are also often associated with poor bone health^3^ and hearing loss due to structural damage to the temporal bone^4,5^. In addition to classical mitochondrial diseases, conditions where heteroplasmy is prevalent include aging, cancer, and neurodegenerative and metabolic diseases. However, studying mitochondrial heteroplasmy in mammalian cells to identify the pathological mechanisms underlying these varied diseases is highly challenging^6^ as it requires the genetic modification of mitochondrial DNA, which has not been achieved *in vivo*.

To overcome this, a heteroplasmic mouse model was developed on the C57BL/6 background, carrying a variable mixture of two mtDNA variants, C57 and NZB. NZB mtDNA differs from C57 mtDNA by several mitochondrial single nucleotide polymorphisms (SNPs), including 12 missense mutations, 4 tRNA mutations, 8 rRNA mutations, and 10 mutations in non-coding regions, making the NZB variant more efficient at oxidative phosphorylation^7^. When both variants coexist, cells selectively retain the preferred mitochondria, leading to varying degrees of heteroplasmy across different tissues and cell types^8^. This mitochondrial selection observed in mice closely reflects the mutant load distribution seen in tissues affected by primary mitochondrial diseases^9^. Importantly, heteroplasmic mice also present bone-related alterations, including elongated tibias and femurs, and early-onset kyphosis^10^. These skeletal manifestations, together with the previously described systemic metabolic dysfunction^8,10,11^, make this model compelling to dissect how mitochondrial heteroplasmy influences the fundamental developmental process of osteoclast differentiation, essential for bone homeostasis.

Bone health relies on continuous tissue remodeling via a network of bone matrix synthesizing osteoblasts (OBs) and bone resorbing osteoclasts (OCs). OCs are terminally differentiated multinucleated cells originating from myeloid progenitors in the bone marrow. The demanding environmental conditions during osteoclastogenesis require robust metabolic activity to support differentiation and function^12^. Disruptions in energy metabolism during OC differentiation cause an imbalance in skeletal homeostasis, leading to the development of bone related diseases^12^. Although the role of biogenesis of mitochondria in OC is well established^13,14,15,16,17^, the contribution of other mitochondrial quality control mechanisms remains unclear. In addition to generating new mitochondria, maintaining a functional mitochondrial pool during osteoclast differentiation may also require the selective removal of damaged or less efficient organelles via mitophagy. How mitochondrial adaptation mechanisms contribute to metabolic fine tuning during osteoclast differentiation remains poorly understood, particularly in the context of mtDNA mutations. Interestingly, bone tissue generally exhibits low levels of mtDNA heteroplasmy^9^, suggesting that selective pressure against dysfunctional mitochondria may be particularly stringent in bone-residing cells. Whether mitochondrial selection plays an active role in ensuring successful osteoclast differentiation under heteroplasmic conditions remains unknown.

Here, we discover that mitochondrial heteroplasmy impacts osteoclastogenesis both *in vivo* and *ex vivo* using bone marrow samples from heteroplasmic mice and peripheral blood mononuclear cells (PBMCs) from m.3243A>G patients. In mice, we demonstrate that mtDNA heteroplasmy leads to increased bone density, which we attribute to defective osteoclast formation and function. Mechanistically, we identify a previously unrecognized sequential import of nuclear encoded complex I subunits into mitochondria during early differentiation of OCs, a process that is disrupted in heteroplasmic cells and leads to reduced oxidative phosphorylation and ATP production. In parallel, we reveal that mitochondrial DNA selection is essential for OC maturation, a process in which mtDNA segregation has not been previously explored. Importantly, we demonstrate for the first time that the autophagy inducing metabolite spermidine treatment restores mitochondrial quality and rescues osteoclast differentiation and function in both heteroplasmic mice and cells derived from mitochondrial disease patients. These findings offer a promising strategy for treating mitochondrial disease-associated bone pathologies.

## Results

### Osteoclastogenesis is impaired in heteroplasmic cells

Bone abnormalities have been reported in individuals with primary mitochondrial diseases, particularly those with heteroplasmic mitochondrial mutations^18^. To investigate whether this is caused by defective osteoclast formation, we collected blood from middle-aged individuals with MELAS carrying both healthy and m.3243A>G mutated mtDNA within their cytoplasm. This mutation has been linked previously to bone abnormalities^19^. We first verified systemic bone deficiency in our patient cohort by confirming reduced serum levels of the bone resorption marker, carboxy-terminal cross-linked telopeptide of type 1 collagen (CTX-1) (Extended Data Fig. 1a). Next, using peripheral blood mononuclear cells (PBMCs) from age- and sex-matched healthy donors and people with m.3243A>G heteroplasmy, we established an *in vitro* OC differentiation assay^20^. By day 6, mature OCs were fixed for tartrate-resistant acid phosphatase (TRAP) analysis. Using this assay, we found that the formation of mature OCs was significantly reduced in m.3243A>G cells compared to healthy cells (Fig. 1a).

**Figure 1.**
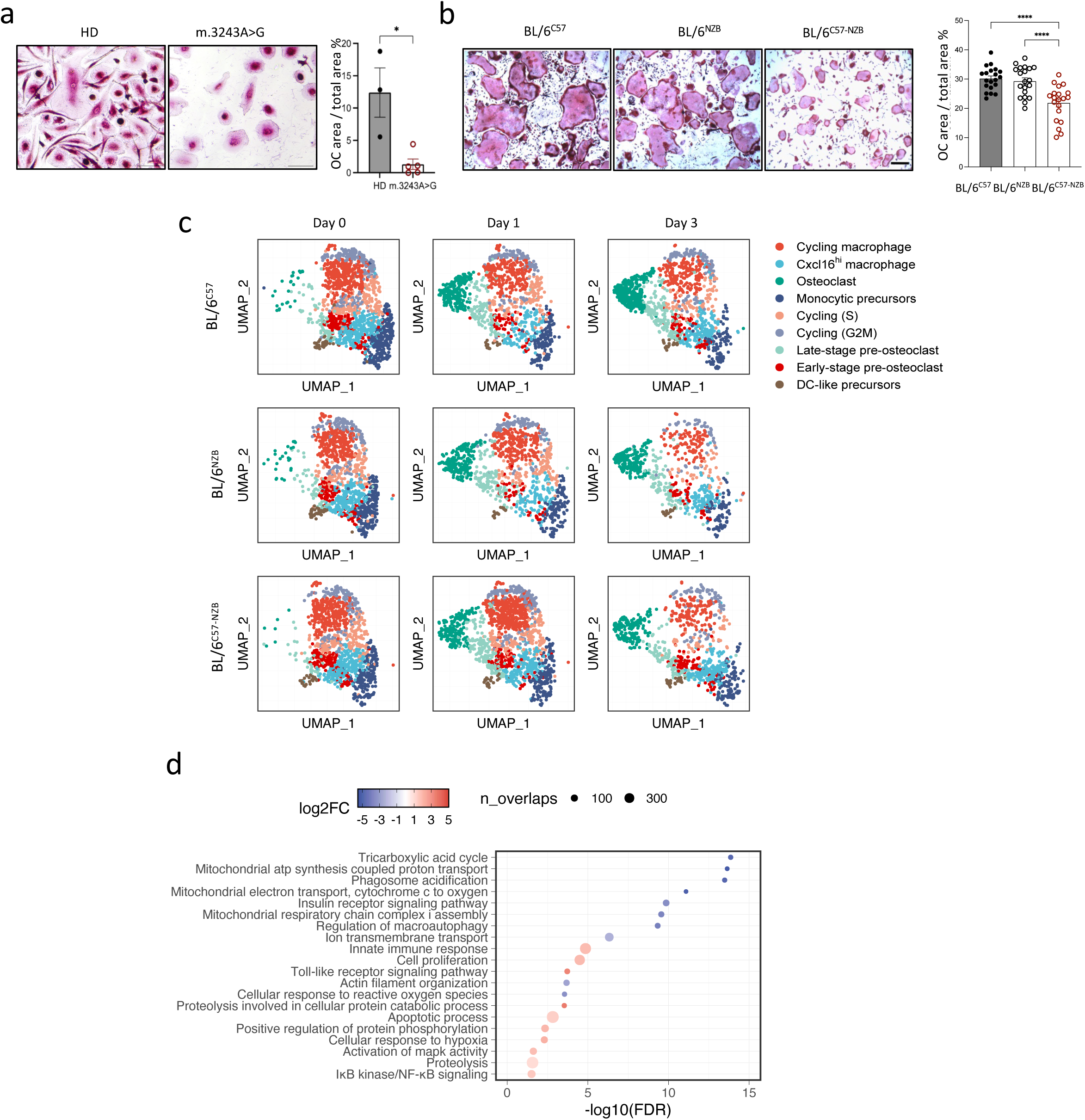
Impaired OC differentiation in heteroplasmic cells, with maturation block emerging within the first three days. **a** Human PBMCs were seeded and differentiated into OCs using 50 ng/ml human M-CSF and 75 ng/ml human RANKL. Left: Representative images of TRAP staining from OC on day 9. Right: Quantification of the percentage of OC area relative to the total area. n=3 healthy donors (HD) and n=5 m.3243A>G patients; 3 technical replicates per condition. * p < 0.05, two-tailed unpaired t-test. Scale bar represents 100 μm. **b** Bone marrow cells from 8-week-old mice BL/6^C57^, BL/6^NZB^ and BL/6^C57-NZB^ mice were differentiated into OCs using 10 ng/ml M-CSF and 50 ng/ml RANKL. OCs were identified by TRAP staining on day 5 (left), scale bar: 200 μm. The percentage of OC area relative to the total area was quantified (right). n= 20 mice per genotype (four independent experiments; each dot represents and individual animal). ****p < 0.0001 determined by one-way ANOVA test. Data are presented as mean ± SEM. **c** UMAP visualization of 10,809 cells during OC differentiation, clustered by transcriptomic similarity and shown at day 0, day 1, and day 3 for different mtDNA groups (BL/6^C57^, BL/6^NZB^, and BL/6^C57-NZB^). **d** GO-BP pathway enrichment analysis for genes upregulated (red) or downregulated (blue) in BL/6^C57-NZB^ OCs compared to BL/6^C57^ OCs.

As introducing mitochondrial point mutations in vivo is challenging, we used the heteroplasmic mouse model (BL/6^C57-NZB^), a unique and well characterized tool in the field^8,10, 21^ to investigate the molecular mechanisms by which mitochondrial heteroplasmy impairs osteoclast differentiation. Bone marrow cells from BL/6^C57-NZB^ heteroplasmic mice, and from BL/6^C57^ and BL/6^NZB^ homoplasmic controls, were cultured *in vitro* for 2 days with murine M-CSF to generate macrophages (day 0), followed by culture with murine M-CSF and RANKL for 5 days to induce OC formation. Consistent with our findings in human cells, bone marrow cells from BL/6^C57-NZB^ mice showed a significant reduction in TRAP-positive OC formation compared to homoplasmic controls (Fig. 1b). These results suggest that it is heteroplasmy itself, rather than complete mitochondrial and nuclear genome mismatch as in the BL/6^NZB^ cells, that underlies the impaired OC differentiation observed in both the BL/6^C57-NZB^ cells and patient samples carrying the m.3243A>G mutation.

To understand the molecular mechanisms underlying this reduced osteoclastogenic capacity, we conducted single-cell RNA sequencing (scRNA-seq) at various stages of OC differentiation (Extended Data Fig. 1b). Bone marrow cells from the heteroplasmic and homoplasmic strains were cultured for 2 days with M-CSF to generate macrophages (day 0), followed by culture with M-CSF and RANKL for either 1 day or 3 days to induce OC formation. Successful *in vitro* OC differentiation was validated by cross-referencing reported marker genes (e.g. *Acp5* and *Ctsk* in Extended Data Fig. 1c) and negative regulators of OC following RANKL treatment (e.g. *Irf8* and *Ma@*)^1^ (Extended Data Fig. 1d). After applying quality control filters, we analyzed a total of 10,809 cells (Extended Data Fig. 1e). Unsupervised graph-based clustering of these single-cell transcriptomic profiles defined 9 distinct cell populations, including a monocyte-like precursor population and a terminal OC population. Other transitional populations included *Cxcl16*^hi^ macrophages, dendritic cells, early- and late-stage pre-OCs, and three proliferative populations (Fig. 1c and Extended Data Fig. 1e).

To infer the linkage between each cell population, we computed a per-cell latent time and performed partition-based graph abstraction (PAGA) analysis based on the RNA-velocity dynamics^22,23^(Extended Data Fig. 1f and g). These analyses revealed a cell trajectory during osteoclastogenesis, which progresses from monocytic precursors to OCs, passing through *Cxcl16*^hi^ macrophages, early-stage pre-OCs and late-stage pre-OCs (Extended Data Fig. 1g). Notably, BL/6^C57-NZB^ cells were relatively concentrated in Cxcl16^hi^ macrophages (17.4%), and early-stage pre-OCs (8.0%) compared to wild-type BL/6^C57^ (14.0% and 6.5%) or BL/6^NZB^ (14.3% and 6.5%) cells. Consequently, fewer heteroplasmic cells reached the mature OC population (10.2% compared to 14.7% and 14.9% in BL/6^C57^ and BL/6^NZB^ cells, respectively) (Fig. 1c and Extended Data Fig. 1g and h).

Given the high metabolic demands and finely tuned cellular adaptive mechanisms of osteoclastogenesis^26^, we hypothesized that mitochondrial heteroplasmy disrupts the coordination of bioenergetic pathways necessary for full osteoclast maturation. To test this, we performed pathway enrichment analysis, which revealed key differences in cellular respiration between the genotypes, including downregulation of the tricarboxylic acid cycle and of mitochondria-related functional pathways in differentiated OCs (Fig. 1d). Despite the well-established hypoxic and oxidative stress environment^24,25^, coupled with the high anabolic demands characteristic of osteoclastogenesis^26^, we found that genes involved in reactive oxygen species handling were unexpectedly downregulated in BL/6^C57-NZB^ cells, accompanied by a shift toward catabolic metabolism (Fig. 1d).

Collectively, our findings suggest that heteroplasmic cells face challenges in osteoclastogenesis, leading to premature arrest at the macrophage stage, which may be linked to altered metabolic adaptation and pathway dysregulation.

### Alterations in complex I in heteroplasmic cells impair metabolic adaptation and oxidative phosphorylation during OC differentiation

We next analyzed the expression of genes involved in major metabolic pathways throughout osteoclast differentiation in these strains. At day 0 of *in vitro* OC differentiation, there were no significant differences in metabolic pathways of BL/6^C57-NZB^ cells compared to wild-type BL/6^C57^; only few pathways, including glycolysis, were upregulated at day 1 upon RANKL treatment in BL/6^C57-NZB^ cells (Fig. 2a). In contrast, after 3 days of differentiation, most mitochondrial-associated pathways were downregulated in heteroplasmic OCs, including ketogenesis, Krebs cycle, galactose metabolism, pentose phosphate metabolism, pyrimidine biosynthesis, adenosine biosynthesis, and glutamate degradation pathway (Fig. 2a).

**Figure 2.**
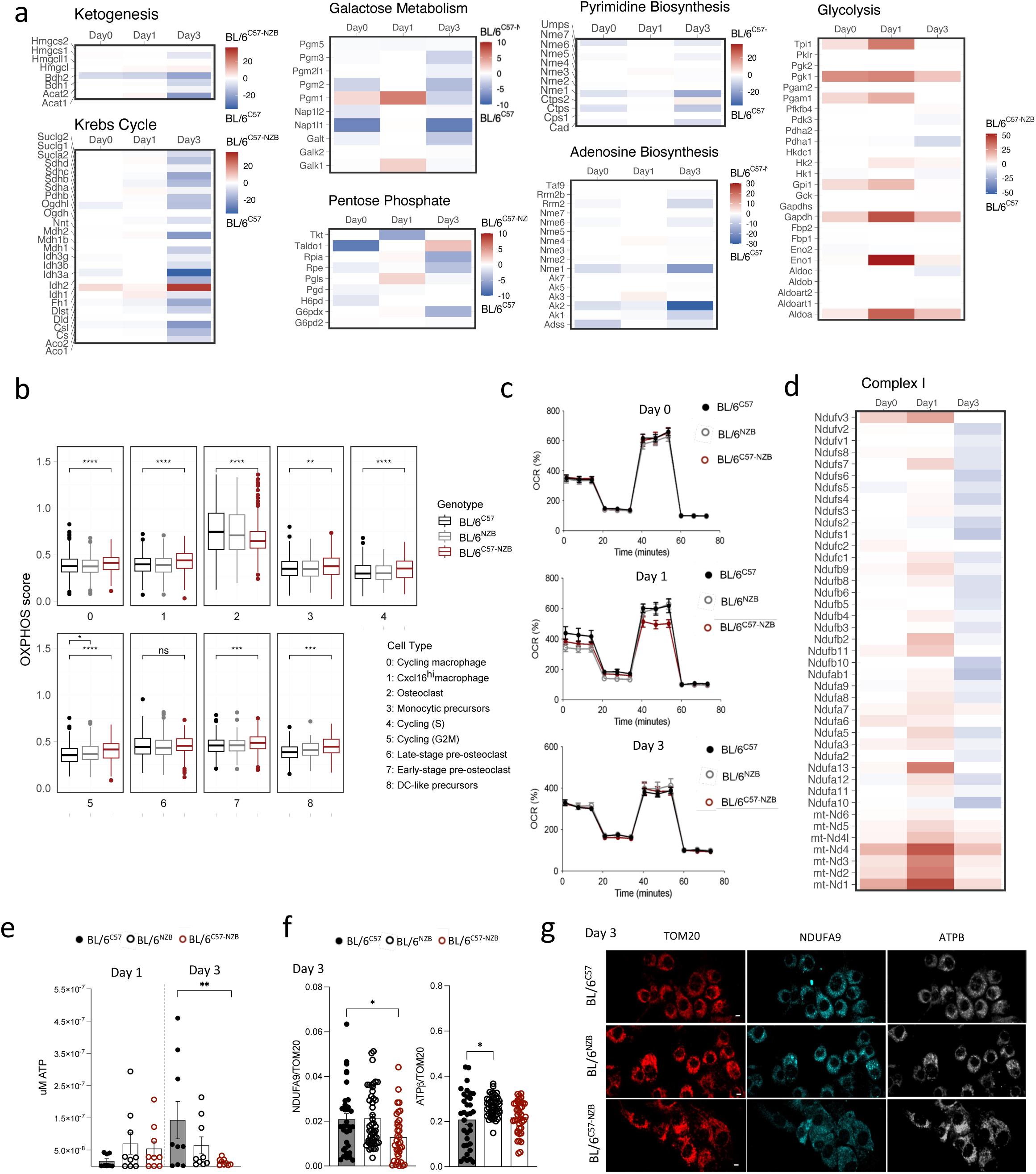
OC profiling reveals metabolic and mitochondrial dysregulation during differentiation in heteroplasmic cells. **a** Heatmap showing gene expression changes in genes encoding subunits involved in ketogenesis, Krebs cycle, galactose metabolism, pentose phosphate, pyrimidine biosynthesis, adenosine biosynthesis and glycolysis in OCs derived from BL/6^C57-NZB^ compared to BL/6^C57^ mice at day 0, day 1, and day 3 (n=3 mice per genotype). **b** Aggregated expression of OXPHOS genes (OXPHOS score) of *in vitro* OCs from different clusters (1-8) derived from BL/6^C57^, BL/6^NZB^, and BL/^6C57-NZB^ mice (n=3 animals per genotype). Module expression scores were normalized by subtracting the aggregated expression levels of a randomly selected, expression matched control gene set. **c** Oxygen consumption rate (OCR) profile of intact cells measured by Seahorse at days 0, 1, and 3 of OC differentiation (3-5 independent assays per biological replicate). Data are presented as mean values ± SEM. **d** Heatmap showing gene expression changes for genes encoding respiratory Complex I in cells derived from BL/6^C57-NZB^ compared to BL/6^C57^ mice at day 0, day 1, and day 3 of OC differentiation (n=3 mice per genotype). **e** Total ATP levels measured by luminometry on days 1 and 3 of osteoclast differentiation in cells from BL/6^C57^, BL/6^NZB^, and BL/6^C57-NZB^ mice. Each dot represents an individual mouse of the indicated genotypes. ** p < 0.01, determined using Tukey’s multiple comparisons following two-way ANOVA. **f-g** Confocal imaging of mitochondrial TOM20, Complex I subunit NDUFA9 and Complex V subunit ATPβ in pre-OCs on day 3 of differentiation derived from BL/6^C57^, BL/6^NZB^, and BL/6^C57-NZB^ mice. **f**, Protein expression quantification ratio of NDUFA9 and ATPβ per mitochondrion (NDUFA9/TOM20; ATPβ/TOM20), based on marker signals (Integrated Density, IntDen) on day 3 (n=4 mice per genotype). **g**, Representative confocal images for the indicated markers. Scale bar: 5 μm, n = 4 mice per genotype. * p <0.05, two-tailed Dunnett’s test following one-way ANOVA. Data are presented as mean values ± SEM.

To systemically evaluate the cellular energy needs in individual cells, we aggregated the expression of genes (n = 107, as annotated in the KEGG database) involved in the oxidative phosphorylation (OXPHOS) pathway (Fig. 2b). As expected, OCs (cluster 2, representing the most mature osteoclastic cells) displayed high OXPHOS scores than other cells along the trajectory (Fig. 2b). In contrast, a significant reduction in OXPHOS score was observed in BL/6^C57-NZB^ OCs compared to homoplasmic controls (Fig. 2b).

MtDNA encodes 13 essential polypeptides that form core components of complexes I, III, IV, and V^28^. We noted that all mitochondrial genome-encoded transcripts were expressed at a higher level at day 3 of wild-type OC differentiation (Extended Data Fig. 2a), suggesting increased energy demand in later differentiation stages. To assess the oxygen consumption rate (OCR) in the OCs directly, we performed Seahorse metabolic assays. Consistent with the gene expression data, no differences were observed on day 0. However, by day 1 of OC differentiation, cells from BL/6^C57-NZB^ mice exhibited a reduction in maximal mitochondrial respiratory capacity (SRC) alongside an increase in mitochondrial mass and reactive oxygen species (ROS), likely as a compensatory response (Fig. 2c and Extended Data Fig. 2b - d). These results provide further evidence of compromised metabolic capacity in heteroplasmic cells early in the OC differentiation process. Notably, by day 3, heteroplasmic cells exhibited lower mitochondrial mass while maintaining the same spare respiratory capacity (SRC) (Extended Data Fig. 2b and c), likely by utilizing alternative substrates and relying on complex II (succinate dehydrogenase)^27^.

BL/6^C57^, BL/6^NZB^, and BL/6^C57-NZB^ cells share an identical nuclear genome but differ in their mtDNA. Both mitochondrial and nuclear genomes encode subunits of the mitochondrial electron transport chain (mtETC), requiring tight coordination between mitochondrial and nuclear gene expression to ensure proper assembly and function. In the context of heteroplasmy, where multiple mitochondrial genomes coexist, this coordination can be disrupted, potentially impairing the stoichiometric assembly of the mtETC and compromising oxidative phosphorylation. The reduction in bioenergetic capacity in heteroplasmic OCs was further confirmed by analyzing the protein level of key mitochondrial electron transport chain (mtETC) subunits using Western blot (Extended Data Fig 2e). BL/6^C57-NZB^ cells displayed a distinct expression profile, with a delayed and less pronounced increase in NDUFB8 (complex I) after day 3 and only a modest rise in SDHB (complex II) levels (Extended Data Fig. 2e), supporting our findings of reduced complex I representation (Fig. 2d). In contrast, wild-type BL/6^C57^ cells showed a coordinated temporal upregulation of OXPHOS components. NDUFB8 and SDHB increased sequentially and peaked at day 3, followed by a marked rise in ATP5A (complex V) at day 5. UQCR2 (complex III) and MTCOI (complex IV) also rose moderately from day 3 onward. Interestingly, BL/6^NZB^ cells followed a similar pattern to BL/6^C57^, with a sharper SDHB peak and a milder ATP5A increase. While the differences between BL/6^C57^ and BL/6^NZB^ may reflect mitochondrial-nuclear co-adaptation in homoplasmic cells, the divergence observed in BL/6^C57-NZB^ cells highlights the functional impact of mitochondrial heteroplasmy on respiratory complex dynamics during differentiation. These results suggests that even if some heteroplasmic cells managed to differentiate into osteoclasts, they may not have the same mitochondrial metabolic performance.

Complexes I and II both feed electrons into the mtETC but differ in substrates and regulation, making their balance essential for energy production and redox homeostasis^29^. In heteroplasmic cells, this balance was altered. We observed a shift in respiratory complex expression, with reduced levels of complex I (NDUFB8) relative to complex II (SDHB). These differences resulted in a notable drop in the relative abundance and activation of complex I compared to complex II proteins. To determine whether this imbalance translated into functional consequences, we next measured complex activity and found that the altered expression profile correlated with a significantly increased complex I/II activity ratio in BL/6^C57-NZB^ cells specifically at day 3 (Extended Data Fig. 2f).

Since complexes I and II drive ATP production via the respiratory chain, we examined whether heteroplasmy-related expression and activity changes affected ATP levels during osteoclast differentiation. We found that BL/6^C57-NZB^ cells exhibited reduced efficiency in ATP production compared to wild type BL/6^C57^ OCs (Fig. 2e). This analysis examined changes in expression on a per cell basis. Interestingly, when analyzed per mitochondria, heteroplasmic cells showed lower levels of NDUFA9, a nuclear DNA (nDNA)-encoded subunit of respiratory complex I, compared to BL/6^C57^ and BL/6^NZB^, after day 3 of *in vitro* OC differentiation (Fig. 2f and g and Extended Data Fig. 2g). In contrast, ATPβ, a subunit of Complex V (ATPase) responsible for ATP synthesis, showed no difference in expression per mitochondrion in heteroplasmic cells on day 3 (Fig. 2f and g). Altogether, the data suggest that dysregulation specifically of complex I is the primary factor affected in OC differentiation by heteroplasmy.

### OC differentiation requires sequential import of complex I subunits

Nuclear-encoded mitochondrial proteins are synthesized by cytosolic ribosomes as precursors and need to be imported into mitochondria^30^. Among them, the complex I subunit NDUFA9 harbors an N-terminal cleavable pre-sequence that facilitates its mitochondrial targeting^31^. We used confocal imaging to track the localization of the complex I subunit NDUFA9 during osteoclast differentiation. Since this had not been previously examined in wild-type cells, our initial aim was to characterize the sequential import dynamics under physiological conditions. In BL/6^C57^ cells, a distinctive feature of early differentiation was the prominent cytoplasmic presence of NDUFA9 (Fig. 2g and Extended Data Fig. 2g), which progressively shifted to a mitochondrial localization (identified by TOM20 staining) by day 5, indicating efficient import during normal osteoclastogenesis (Fig. 3a, b).

**Figure 3.**
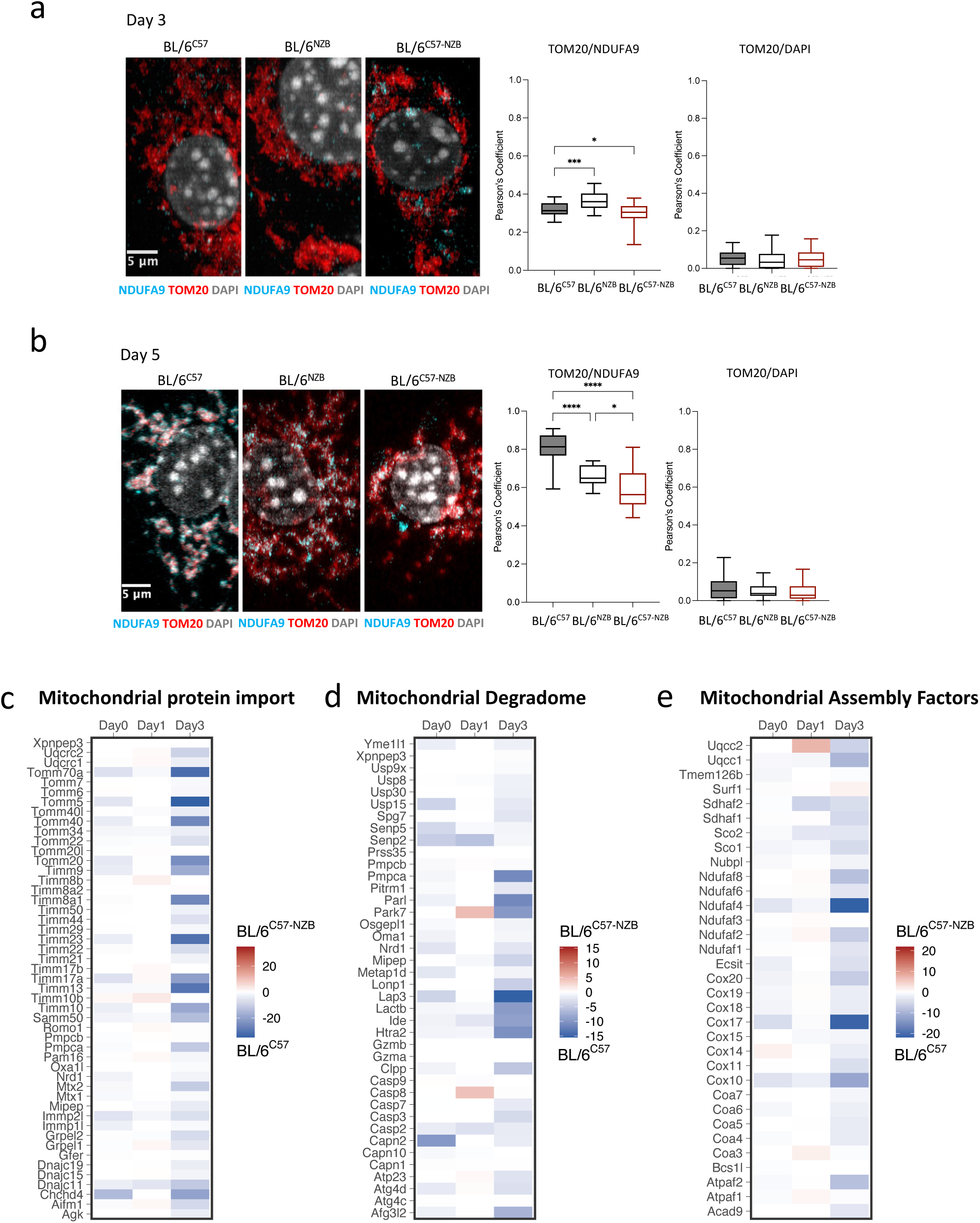
Impaired import of nuclear-encoded complex I subunits disrupts complex assembly and function in heteroplasmic cells. **a-b**: Confocal imaging of mitochondrial TOM20, Complex I subunit NDUFA9 and DAPI in pre-OCs on day 3 of differentiation (**a**) and mature OCs on day 5 of differentiation (**b**) derived from BL/6^C57^, BL/6^NZB^, and BL/6^C57-NZB^ mice. Left: Representative confocal images for the indicated markers. Right: Quantification of colocalization of Complex I in the mitochondria (TOM20/NDUFA9) using as a control colocalization between mitochondria and the nucleus (TOM20/DAPI), shown by Pearson’s coefficient. Scale bar: 5 μm, n = 4 mice per genotype. * p <0.05, *** p < 0.001, **** p < 0.00001, two-tailed Dunnett’s test following one-way ANOVA. In **a-b**, data are presented as mean values ± SEM. **c-e** Heatmap showing gene expression changes for genes involved in the mitochondrial protein import machinery (**c**), mitochondrial degradome (**d**) and mitochondrial assembly factors (**e**) in OCs cultures derived from BL/6^C57-NZB^ and BL/6^C57^ mice at day 0, day 1, and day 3 (n=3 mice per genotype).

Previous studies indicate that complex I alterations disrupts mitochondrial protein import, impairs chaperone function, and activates the mitochondrial unfolded protein response to prevent aggregation of respiratory complex subunits^32^. Consistent with this, we found that the progressive mitochondrial import of NDUFA9 observed in wild-type BL/6^C57^ cells was notably reduced in BL/6^NZB^ cells, and even more markedly in BL/6^C57-NZB^ cells. By day 5, mitochondrial colocalization of NDUFA9 was significantly diminished in both genotypes, suggesting impaired import or retention of this subunit under conditions of heteroplasmy and mitonuclear mismatch (Fig. 3a, b). This suggests that the reduced import of nuclear-encoded complex I subunits arises from impaired mitochondrial import machinery, likely triggered by nucleo-mitochondrial imbalance or stress responses. Supporting this, scRNA-seq revealed reduced nuclear-encoded complex I subunits and increased mitochondrial-encoded genes in heteroplasmic OCs (Fig. 2d), consistent with a stress response that may impair import and complex I assembly.

To directly address this, we analyzed scRNA-seq data for gene expression changes related to mitochondrial protein import and the mitochondrial degradome at days 0, 1, and 3 of OC differentiation. We found pronounced downregulation of genes encoding key components of the mitochondrial protein import machinery in BL/6^C57-NZB^ cells compared to wild type BL/6^C57^ cells (Fig. 3c). Genes involved in mitochondrial protein degradation were also reduced in heteroplasmic cells (Fig. 3d), potentially contributing to the cytoplasmic accumulation of complex I subunits. Furthermore, our data indicates a downregulation of the assembly proteins for respiratory complex I (Fig 3e), suggesting that the failure to complete protein import into mitochondria leads to decreased respiratory complex assembly factors in BL/6^C57-NZB^ cells compared to BL/6^C57^.

Altogether, our data suggest that nuclear-encoded complex I subunits fail to be either imported into mitochondria or properly degraded in the cytoplasm during OC differentiation in BL/6^C57-NZB^. Both processes require ATP^30,33,34^, which is reduced with heteroplasmy at the stage when OCs would normally depend on respiratory complex I formation for mitochondrial function.

### Mitochondrial selection in heteroplasmic cells is essential for osteoclast differentiation

We previously reported that mitochondrial heteroplasmy can be resolved through non-random mtDNA selection, a process that is tissue- and cell-type specific^8^. Given the substantial metabolic shifts that occur during osteoclastogenesis^35^, we hypothesized that different stages of OC differentiation impose distinct bioenergetic demands, potentially requiring active mitochondrial selection.

To evaluate mtDNA proportion changes during differentiation, we adapted the mgatk pipeline to genotype mitochondrial variants at single-cell resolution using deep scRNA-seq libraries during early OC differentiation^36^. After strict filtering, we were able to recover on average 237 reads per cell covering the NZB mtDNA haplotype (Extended Data Fig. 3a). We observed a higher level of NZB mtDNA transcripts on day 3 compared to day 1 cells, independent of the sequencing depth (Fig. 4a). The percentage of NZB mtDNA also slightly increased along the OC differentiation trajectory (Extended Data Fig. 3b). We further validated these findings using PCR followed by RFLP (Restriction Fragment Length Polymorphism) analysis^21,37,38^ in OC cultures at Days 1 and 3. Our data showed a shift toward more NZB mtDNA as OC differentiation progresses (Fig. 4b and Extended Data Fig. 3c, d). This change in mtDNA proportion from day 1 to day 3 was driven by the formation of mature OCs, which carried increased NZB mtDNA content (Fig. 4c). Other cells did not show significant enrichment for the NZB mtDNA haplotype compared to monocytic precursors (Fig. 4c). These results suggest that the selection of the preferred mtDNA variant, reflected in the higher proportion of NZB mtDNA in mature OCs, facilitates OC maturation. Moreover, the selection towards a mtDNA haplotype different from the nuclear genome suggests that the pathogenic phenotypes observed in heteroplasmic OCs were indeed caused by heteroplasmy per se, rather than specific SNPs from the NZB mitochondrial genome.

**Figure 4.**
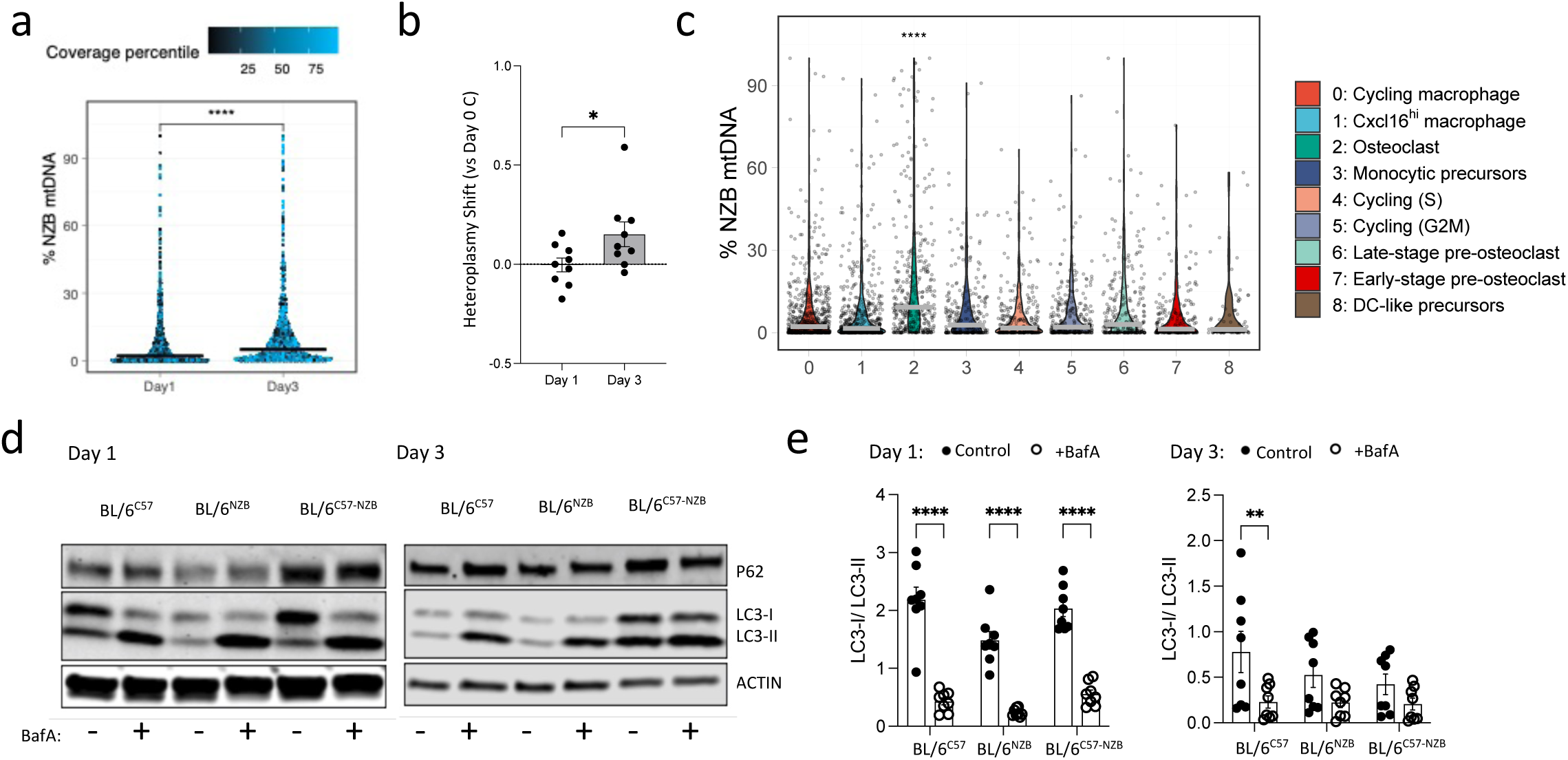
OC differentiation relies on sustained autophagy during early stages, with mtDNA selection required in heteroplasmic cells. **a** NZB mtDNA levels (estimated as a percentage of total mtDNA) from BL/6^C57-NZB^ mice, showing a shift toward the NZB haplotype in day 3 cells compared to day 1. Each dot represents one cell and is colored by the percentile-normalized sequencing coverage of NZB mtDNA variants. The color scale represents sequencing depth. Day 1: n = 1,249 cells. Day 3: n = 884 cells. Median values are marked by black lines. **** p < 0.0001, Wilcoxon test. **b** Assessment of the heteroplasmy shift toward NZB mtDNA at day 3 compared to day 1 in pre-OCs derived from BL/6^C57-NZB^ young mice (n=9; each dot represents an individual mouse). The transformed heteroplasmy shift was evaluated using day 0 as the reference for each heteroplasmic mouse. NZB mtDNA levels were obtained by PCR, followed by RFLP (Restriction Fragments Length Polymorphism) to distinguish between C57 and NZB mtDNA variants. * p < 0.05, unpaired t test. Data are presented as mean values ± SEM. **c** NZB mtDNA levels in BL/6^C57-NZB^ cells across different clusters indicating increased NZB mtDNA selection in OCs compared to monocytic precursors. Median values are marked by grey lines. **** p < 0.0001, Wilcoxon test, with monocytic precursors as the reference group. **d-e** Western Blot analysis of autophagy in primary murine BM cells treated with M-CSF and RANKL for 1 or 3 days. **d,** Representative images of membranes showing p62 and LC-3 expression. **e,** autophagy flux quantification of the LC-3 bands (LC3-I/LC3-II) at day 1 and 3 of OC differentiation. 10 nM BafA1 was added 2 hours before protein harvest, with actin used as a protein loading control. n = 8 independent biological samples for DMSO control and BafA. **p < 0.01, ****p < 0.0001, two-way ANOVA with Sidak’s multiple comparisons test. Data are presented as mean values ± SEM.

While failed mitochondrial selection has previously been associated with impaired autophagy^10^, it remains unclear whether mitophagy clears unwanted mitochondria in mtDNA heteroplasmic osteoclasts. On day 1, BafA1 treatment increased LC3-II levels and decreased the LC3-I/LC3-II ratio, indicating a strong autophagic flux across all genotypes (Fig. 4d, e). However, by day 3, BafA1, which was used to block autophagosome-lysosome fusion and assess autophagy flux, no longer increased LC3-II in BL/6^C57-NZB^ cells (Fig. 4d, e). This, together with elevated p62 levels in heteroplasmic cells, even in the absence of BafA1 (Fig. 4d and Extended Data Fig. 3e), suggests a block in autophagic flux at day 3. Together, these results suggest that reduced autophagy throughout differentiation may underlie the metabolic alterations observed in heteroplasmic OCs.

### Osteoclast maturation in murine and human heteroplasmic cells can be restored by promoting mtDNA selection through spermidine supplementation

We previously demonstrated that autophagy is regulated by endogenous spermidine levels which control translation of TFEB^39,40^. Spermidine also activates mitochondrial translation in myeloid cells^41^ and boosts oxidative metabolism by binding to the mitochondrial trifunctional protein (MTP) in T cells^42^. Spermidine given to mice in the drinking water restores autophagy levels when these are low^43^. Therefore, we used spermidine to directly test whether it improves mitochondrial function and resolves the altered metabolic phenotype observed in heteroplasmic OCs. First, we performed OC differentiation of cells from heteroplasmic mice with spermidine supplementation in the culture media. Spermidine restored OC formation in heteroplasmic cells *in vitro* compared to control cultures (Fig. 5a, b). Interestingly, the spermidine dose required to rescue OC differentiation in heteroplasmic cells has previously been shown to inhibit osteoclast formation in wild-type cells^44^ - a result we also confirmed in our analyses (Fig. 5a, b). This suggests that heteroplasmic cells have exhausted their endogenous spermidine levels and now have a higher demand for spermidine to support proper differentiation. Spermidine supplementation significantly increased NZB mtDNA selection on day 5 (Fig. 5c), while the total mtDNA content remained stable (Extended Data Fig. 4a).

**Figure 5.**
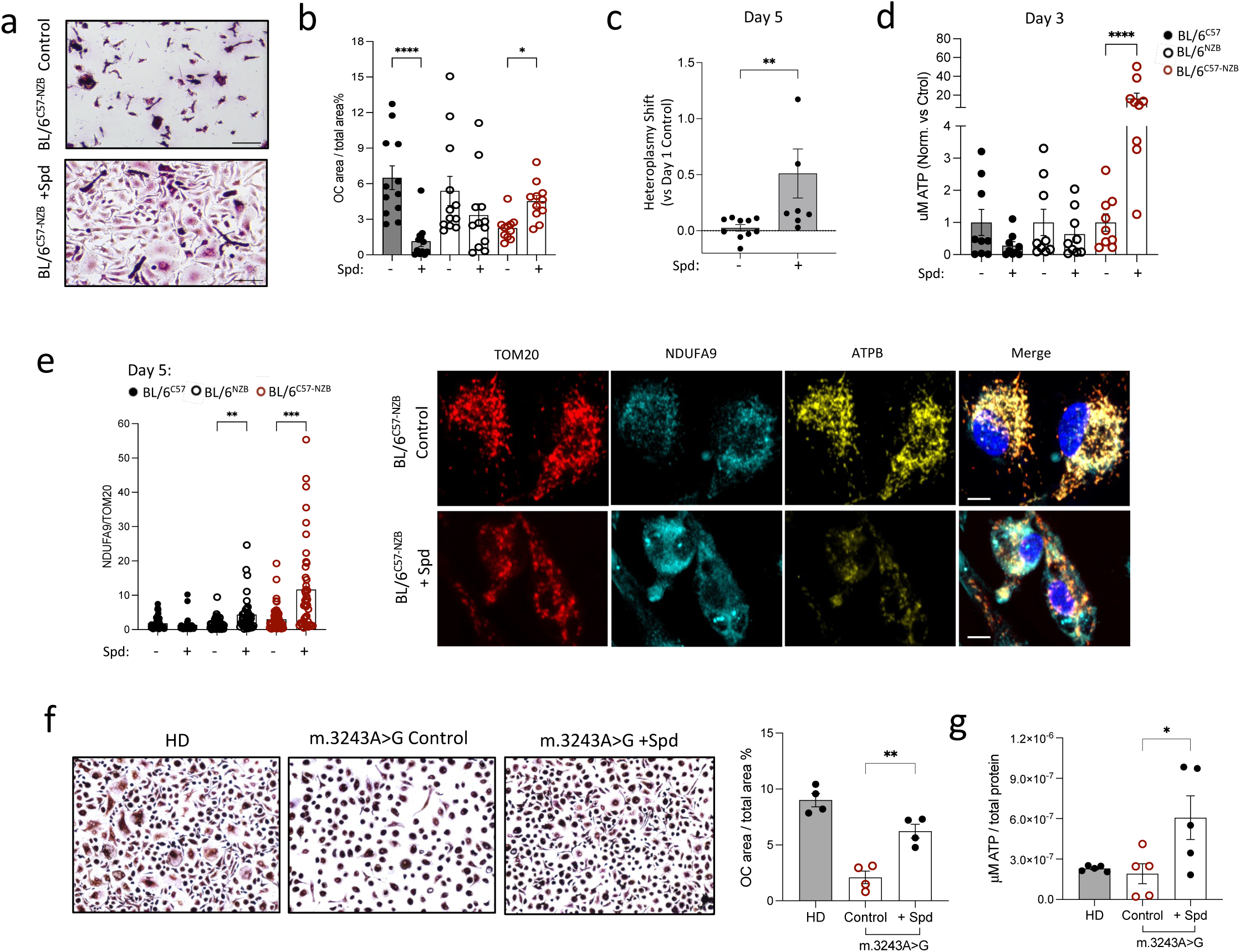
Spermidine enhances cellular ATP levels and restores osteoclast differentiation in murine and human heteroplasmic cells. **a-b** Bone marrow cells from 8-week-old BL/6^C57^, BL/6^NZB^ and BL/6^C57-NZB^ mice were differentiated into OCs using 10 ng/ml M-CSF and 50 ng/ml RANKL with or without 10 uM spermidine (spd) treatment. **a,** OCs were identified by TRAP staining on day 5. Scale bar: 50 μm. **b,** percentage of OC area relative to the total area was quantified. Data are representative of two independent experiments. * p < 0.05, **** p < 0.0001, nonparametric Kruskal-Wallis test. Data are presented as mean values ± SEM. **c** Assessment of the heteroplasmy shift toward NZB mtDNA at day 5 in OCs derived from BL/6^C57-NZB^ young mice (n=10 for untreated and n=8 for cultures treated with 10 µM spermidine; each dot represents an individual mouse). The transformed heteroplasmy shift was evaluated using day 1 as the reference for each heteroplasmic mouse. NZB mtDNA levels were quantified by PCR, followed by RFLP (Restriction Fragments Length Polymorphism) to distinguish between C57 and NZB mtDNA variants. ** p < 0.01, nonparametric Mann-Whitney test. Data are presented as mean values ± SEM. **d** Total ATP levels were measured by luminescence on day 3 of OC differentiation in cells from BL/6^C57^, BL/6^NZB^, and BL/6^C57-NZB^ mice, with and without 10 µM spermidine treatment. Data are normalized to untreated cells and each dot represents a sample from an individual mouse of the indicated genotype. **** p < 0.0001, one-way ANOVA and Sidak’s multiple comparison test. Data are presented as mean values ± SEM. **e** Confocal imaging of mitochondrial TOM20, Complex I subunit NDUFA9, and ATPase subunit ATPβ in OCs derived from BL/6^C57^and BL/6^NZB^, and BL/6^C57-NZB^ mice with and without spermidine treatment on day 5 of *in vitro* OC differentiation. Left: Protein quantification ratio of NDUFA9 per mitochondrion (NDUFA9/TOM20), based on marker signals (Integrated Density, IntDen) (n=3 mice per genotype). ** p <0.01, *** p <0.001, two-tailed Dunnett’s test following one-way ANOVA. Data are presented as mean values ± SEM. Right: Representative confocal images for the indicated markers in pre-OCs derived from BL/6^C57-NZB^ mice with and without spermidine treatment on day 5 of *in vitro* OC differentiation. Scale bar: 5 μm. **f** Human PBMCs were seeded and differentiated into OCs in the presence of 50 ng/ml human M-CSF and 75 ng/ml human RANKL, with or without 10 µM spermidine treatment. Left: Representative images of TRAP staining from OC on day 6. Scale bar: 50 μm. Right: Quantification of the percentage of OC area relative to the total area. n=4 healthy donors (HD) and n=4 m.3243A>G patients; 6 technical replicates per condition. ** p < 0.01, determined using Tukey’s multiple comparisons following one-way ANOVA. Data are presented as mean values ± SEM. **g** Total ATP levels assessed by luminescence on day 5 of OC differentiation in cells derived from human PBMCs in the presence of 50 ng/ml human M-CSF and 75 ng/ml human RANKL, with or without 10 µM spermidine treatment. Each dot represents *in vitro* OC differentiation from a donor. n=5 healthy donors (HD) and n=5 m.3243A>G patients. * p < 0.05, determined using Tukey’s multiple comparisons following one-way ANOVA. Data are presented as mean values ± SEM.

While no changes in total ATP were observed in the spermidine-treated OC cultures on day 1 of differentiation (Extended Data Fig. 4b), treated heteroplasmic cells showed a sharp increase in ATP content on day 3 (Fig. 5d). Interestingly, despite this increase, ATPβ levels were reduced in BL/6^C57^, BL/6^NZB^ and BL/6^C57-NZB^ cells by day 5 (Extended Data Fig. 4c, d). Confocal imaging revealed increased mitochondrial complex I expression relative to mitochondrial content specifically in spermidine-treated OC cultures from BL/6^C57-NZB^ mice on day 5, with no similar effect observed in the other genotypes (Fig. 5e and Extended Data Fig. 4d). These findings suggest that spermidine treatment improves mitochondrial efficiency and promotes ATP synthesis by boosting complex I function and mitochondrial respiratory capacity, rather than simply increasing ATP synthase levels.

To extend the relevance of our findings to a human context, we tested whether spermidine could improve osteoclast differentiation in cells derived from individuals carrying the m.3243A>G mitochondrial mutation. While spermidine has been previously linked to autophagy and mitochondrial maintenance, its effects have not been evaluated in the setting of mitochondrial heteroplasmy in primary human cells. Treatment with 10 µM spermidine increased both the number of TRAP⁺ OCs (Fig. 5f) and intracellular ATP levels (Fig. 5g) in m.3243A>G cells, supporting its potential to enhance mitochondrial function and restore differentiation capacity in this pathogenic context.

Together, these results position spermidine as a promising metabolite supplement capable of improving the phenotype of cells that require mitochondrial heteroplasmy resolution for proper differentiation and function.

### BL/6^C57-NZB^ mice exhibit increased bone density due to diminished OC function

Next, we assessed how mitochondrial heterogeneity in OCs and their impaired differentiation affects bone homeostasis in vivo. Unlike in humans, the heteroplasmic mouse model lacks nuclear genetic variation between individuals, allowing us to specifically study the effects of mitochondrial differences without genetic confounders, providing a controlled setting to investigate this phenomenon *in vivo*. We first investigated whether bone-resorbing OCs are affected in BL/6^C57-NZB^ mice by staining longitudinal cross-sections of tibiae with TRAP specific for OC. According to bone histomorphometry, tibiae from BL/6^C57-NZB^ mice had significantly lower signal for TRAP-positive OC when compared to those from BL/6^C57^ control mice (Fig. 6a, b). Consistently, a significant reduction of CTX-1 level was observed in serum from BL/6^C57-NZB^ mice, indicating a lower level of systemic OC activity and bone resorption (Fig. 6c). No differences were observed in the frequency of bone marrow OC precursors as demonstrated by expression of surface markers (CD11b^−^CD3^−^B220^−^CD115^+^) between the 3 genotypes, indicating specifically a diminished OC differentiation from BL/6^C57-NZB^ OC precursors (Extended Data Fig. 5a). In addition, no differences were observed in serum procollagen type I N-terminal propeptide (PINP), an OB-derived marker reflecting bone formation, among the three groups (Fig. 6d).

**Figure 6.**
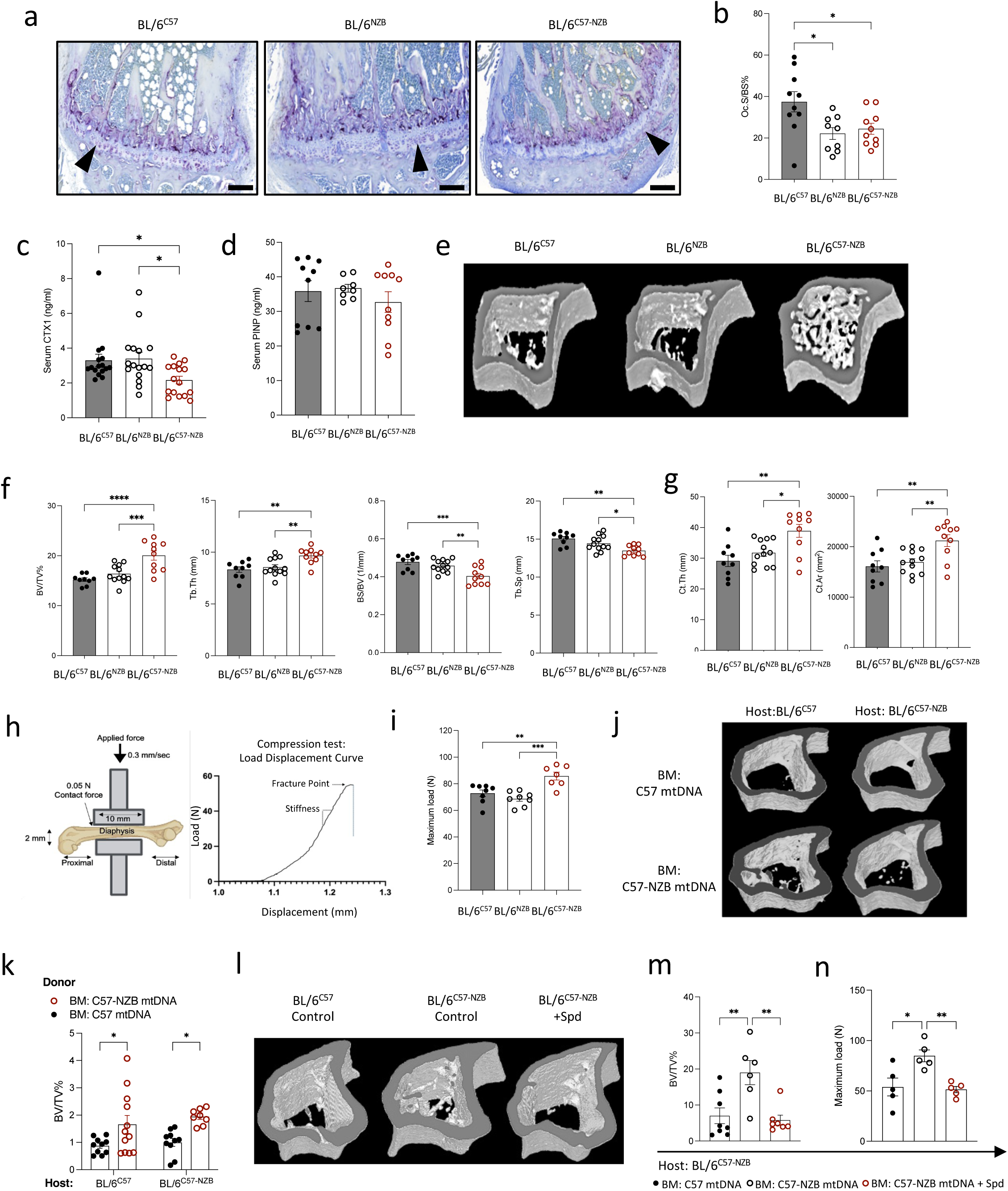
OC dysfunction drives increased bone density in heteroplasmic mice, reversed by spermidine treatment. **a** Representative TRAP/0.2% methyl green-stained tibial sections showing OCs (red/purple, black arrowheads) on the endocortical bone surface in BL/6^C57^, BL/6^NZB^ and BL/6^C57-NZB^ mice (n=5, two independent experiments). Scale bar: 200 μm. **b** Bone histomorphometric quantification of OCs: percentage of OC surface relative to the total bone surface (Oc.S/BS) (BL/6^C57^, *n* = 10; BL/6^NZB^, *n* = 9; and BL/6^C57-NZB^, *n* = 10 mice, each dot represents an individual animal). * p < 0.05, one-way ANOVA followed by Tukey’s multiple comparisons test.Data are presented as mean ± SEM. **c-d** ELISA analysis of serum levels of CTX-1 (**c**) (BL/6^C57^, *n* = 16; BL/6^NZB^, *n* = 16; and BL/6^C57-NZB^, *n* = 16 mice) and PINP (**d**) (BL/6^C57^, *n* = 10; BL/6^NZB^, *n* = 8; and BL/6^C57-NZB^, *n* = 10 mice), each dot represents an individual animal. * p < 0.05, one-way ANOVA followed by Tukey’s multiple comparisons test.Data are presented as mean ± SEM. **e** Representative micro-CT reconstructed images of tibiae from 12-week-old male BL/6^C57^, BL/6^NZB^ and BL/6^C57-NZB^ mice. **f**–**g** Micro-CT analysis of trabecular and cortical bone morphology parameters: (**f**) BV/TV (bone volume fraction), Tb.Th (trabecular thickness), BS/BV (bone surface density), Tb.Sp (trabecular separation); (**g**) Ct.Th (cortical thickness) and Ct.Ar (cortical area) (BL/6^C57^, *n* = 9; BL/6^NZB^, *n* = 12; and BL/6^C57-NZB^, *n* = 10; each dot represents tibial measurements from an individual animal). * p < 0.05, ** p < 0.01, *** p < 0.001, **** p < 0.0001, one-way ANOVA followed by Tukey’s multiple comparisons test. Data are presented as mean ± SEM. **h-i** Schematic overview illustrating the positioning of femur for compression testing between the load area and the platform load frame (**h, left**). Representative load displacement curve of femur from the BL/6^C57^ control group (**h, right**). Dot plot showing the maximum load values for the femur in 12- to 14-week-old males of the indicated genotypes (**i**). BL/6^C57^, *n* = 8; BL/6^NZB^, *n* = 8; and BL/6^C57-NZB^, *n* = 7. Each dot represents femoral measurements from an individual animal. ** p < 0.01, *** p < 0.001, one-way ANOVA followed by Tukey’s multiple comparisons test. Data are presented as mean ± SEM. **j-k** Lethally irradiated 8-week-old CD45.1+ B6.SJL and BL/6^C57-NZB^ CD45.1+ mice were reconstituted with BM cells from BL/6^C57^ CD45.2+ or BL/6^C57-NZB^ CD45.2+ mice. After 9 weeks, micro-CT analysis was performed on tibiae. **j,** Representative micro-CT reconstructed images. **k,** BV/TV quantification (BL/6^C57^ to CD45.1+ B6.SJL, n = 10; BL/6^C57-NZB^ to CD45.1^+^ B6.SJL, n = 12; BL/6^C57^ to BL/6^C57-NZB^, n = 10; BL/6^C57-NZB^ to BL/6^C57-NZB^, n = 8; each dot represents an individual animal). *p < 0.05, two-way ANOVA. Data are presented as mean values ± SEM. **l-m** Lethally irradiated young BL/6^C57-NZB^ mice were reconstituted with bone marrow from BL/6^C57^ or BL/6^C57-NZB^ mice and treated with 5 mM spermidine in drinking water for 8 weeks. **l,** Representative micro-CT reconstructed images of tibiae from the bone marrow reconstituted mice. **m,** BV/TV of tibiae quantification by micro-CT (n = 8 BL/6^C57^; n = 6 BL/6^C57-NZB^ and n = 7 BL/6^C57-NZB^ + Spd; each dot represents an individual animal). The experiment is representative of two independent experiments. ** p < 0.01, one-way ANOVA followed by Tukey’s multiple comparisons test. Data are presented as mean ± SEM. **n** Bending test analysis in bone marrow transplanted mice treated with 5 mM spermidine. Dot plots of maximum femur load from bone marrow-transplanted mice, assessed using the bending test (n = 5 mice per BM cell genotype; each dot represents an independent animal). *p < 0.05, **p < 0.01, one-way ANOVA followed by Tukey’s multiple comparisons test. Data are presented as mean ± SEM.

To investigate if bone mass is affected in BL/6^C57-NZB^ mice, tibiae were collected from aged-matched male BL/6^C57^, BL/6^NZB^ and BL/6^C57-NZB^ mice for micro-CT analysis. BL/6^C57-NZB^ mice showed an increase in trabecular bone (Fig. 6e). Accordingly, quantification of bone parameters revealed a 1.3-fold increase of bone volume over total volume (BV/TV) and a 1.2-fold increase of trabecular thickness (Tb.Th), accompanied by a significant decrease in bone surface over bone volume (BS/BV) and in trabecular separation (Tb.Sp) (Fig. 6f). The decrease in the BS/BV ratio in BL/6^C57-NZB^ group was due to a significant increase in BV, as compared to BS. Furthermore, a significant increase of cortical thickness (Ct.Th) and cortical area (Ct.Ar) was observed in the BL/6^C57-NZB^ group (Fig. 6g), confirming amelioration of overall bone parameters. Next, we investigated whether these changed bone parameters affected bone strength by measuring resistance to fracture. We conducted biomechanical compression testing of the femur from BL/6^C57^ mice, BL/6^NZB^ mice and BL/6^C57-NZB^ mice (Fig. 6h). Compared with femurs from BL/6^C57^ mice, those from BL/6^C57-NZB^ mice needed 25% more maximum load to break (Fig. 6i). Therefore, the femur from BL/6^C57-NZB^ had increased bone mechanical strength, resulting in greater resistance to deformation or fracture.

Our results showing impaired OC differentiation, along with serum CTX-1 and PINP analyses, indicate that the bone alterations observed in BL/6^C57-NZB^ mice are likely driven by diminished OC activity, rather than changes in other non-hematopoietic cells such as OBs. To test this, we performed bone marrow (BM) chimera experiments, where the host provides the OBs and the donor supplies the OCs, given their hematopoietic origin (Fig. 6j, k and Extended Data Fig. 5b-d). Lethally irradiated BL/6^C57-NZB^ and BL/6^C57^ mice were transplanted with BM donor cells from either BL/6^C57^ or BL/6^C57-NZB^ mice. BM engraftments were monitored by bleeding every 2 weeks, with the donor cells expressing the CD45.2 isoform while the host was CD45.1^+^ to track donor and host cells (Extended Data Fig. 5b-c). Notably, we observed an overall increase in bone density, as indicated by BV/TV measurements, when the BM was heteroplasmic (Fig. 6j, k), along with changes in additional parameters assessed by micro-CT analysis (Extended Data Fig. 5d), regardless of the recipient strain. These findings indicate that the increased bone volume in BL/6^C57-NZB^ mice is driven by impaired differentiation and formation specifically of OCs, which are of hematopoietic origin.

Building on our *in vitro* findings that spermidine restores OC differentiation in heteroplasmic cells, we next investigated whether spermidine could also modulate this phenotype *in vivo* (Fig. 6l-n and Extended Data Fig. 5e). We generated BM chimeric mice by transplanting BM donor cells from wild-type BL/6^C57^ or BL/6^C57NZB^ mice into lethally irradiated BL/6^C57-NZB^ recipients, ensuring that all recipient mice share the same host tissue background. After 8 weeks of spermidine treatment (5 mM in drinking water), bone phenotyping analyses using micro-CT demonstrated that spermidine administration in BL/6^C57-NZB^ BM donor cells led to a decrease of BV/TV in mice (Fig. 6m), accompanied by an increase in Tb. Sp (Extended Data Fig. 5e), confirming an activation of bone catabolism. Lastly, spermidine-treated chimeric BL/6^C57-NZB^ mice showed decreased bone mechanical strength, reversing the phenotype to resemble wild-type BM (Fig. 6n).

Taken together, OC differentiation in heteroplasmic cells is linked to a shift in mitochondrial heteroplasmy toward NZB mtDNA variants. By restoring spermidine levels that may have been depleted due to the extra demand during heteroplasmic cell differentiation, osteoclastogenesis was restored, both *in vitro* and *in vivo*.

## Discussion

Mitochondrial heteroplasmy can arise in healthy individuals through the random accumulation of mitochondrial DNA mutations and underlies the symptoms of mitochondrial diseases caused by pathogenic variants. These disorders often affect multiple organs, though the reasons for tissue-specific vulnerability remain unclear. In this study, we demonstrate that mitochondrial heteroplasmy impairs the highly energy-demanding process of osteoclast differentiation, specifically at the stage when mitochondrial respiration becomes essential. Importantly, we could uncover for the first time that when nuclear encoded complex I of the mtETC does not match the mitochondrial encoded mtETC components, its import into the mitochondria and thereby the proper assembly of the mtETC is impaired, affecting ATP metabolism. Heteroplasmy leads to premature termination of osteoclastogenesis and thereby reduced bone resorption. We further demonstrate that autophagic flux gets exhausted over the course of heteroplasmic OC differentiation and spermidine, a metabolite needed for autophagy, partially rescues OC function in both heteroplasmic mice and in heteroplasmic patients’ cells.

The generation of new mitochondrial DNA (mtDNA) and proteins has been linked to OC differentiation and function. Indeed, mature OCs have bigger and larger numbers of mitochondria, elevated mitochondrial oxygen consumption rates and higher levels of the mtETC enzymes^13,17^. Notably, RANKL has been shown to stimulate mitochondrial biogenesis^17,45^. The extensive transcriptional and metabolic reprogramming and intact mitochondrial function are essential for OC differentiation and to demineralize the bone.

The assembly of mitochondrial respiratory complex I in mammals requires the coordinated integration of at least 44 subunits encoded by either nuclear or mitochondrial genomes^46^. The presence of complex I subcomplexes (intermediate state) in complex I-deficient cells (i.e. when the complex is not assembled properly) indicates a sequential assembly process^47,48,31^. This involves the complete import of nuclear-encoded subunits into mitochondria, degradation of excess cytoplasmic material, and ultimately, the proper assembly and superassembly of respiratory complex I in the mitochondrial lumen. Although not all mtDNA mutations directly affect mtETC subunits, such as the m.3243A>G MELAS mutation in a mitochondrial tRNA gene, they can still disrupt mitochondrial protein synthesis or quality control, which in turn can impair mtETC subunit expression and function. Our data suggests that mitochondrial protein subunits, in particular complex I, accumulate as unimported mitochondrial proteins in the cytoplasm during OC differentiation. The absence of Ndufs4, a subunit of complex I in the mtETC encoded by the nuclear genome, has been associated with impaired OC differentiation and function, leading to increased bone density^49^. In addition, it has been reported that ECSIT, a complex I-associated protein, plays a central role in the mitochondrial response to early signaling of RANKL^45^. Osteoclastogenesis is a tightly regulated process that demands ATP not only for biomass generation but also for metabolic adaptation to substrate availability at different stages. This process unfolds in a hypoxic tissue environment^25,50^. Intriguingly, hypoxia-induced mitochondrial stress granules have been identified in C. elegans^51^. Similarly, during aerobic growth, yeast cells were shown to accumulate mitochondrial precursor proteins as MitoStores in the cytoplasm^52^. Hence, one explanation for complex I being retained in the cytoplasm may be that this temporary storage site ensures their rapid import into mitochondria during the later stages of differentiation, particularly under challenging nutritional conditions or hypoxic induction. However, unassembled complex I accumulate in the cytoplasm and are bound for degradation. Previous studies in mammalian cells suggest that stress granules can be cleared either through an autophagy-independent disassembly process or via autophagy-dependent degradation^53,54^. The impaired autophagic flux observed in the latter stages during OC differentiation in heteroplasmic BL/6^C57-NZB^ cells may contribute to the cytoplasmic accumulation of complex I.

Mitochondrial heteroplasmy presents a largely unexplored area that may influence cellular function and bone health. Spermidine has been proposed to enhance autophagy^40,55^, translational activity^43^ and mitochondrial metabolism^42,56,57^. Collectively, these findings highlight spermidine as a promising candidate for improving mitochondrial quality. We and others have found that in both mice and humans a moiety of spermidine is needed to activate the translation factor *eif5a* by hypusination in order to increase autophagic flux and lysosomal activity^43,58^. Additionally, this translation factor is crucial for mitochondrial translation^41^, and in T cells boosts oxidative metabolism by binding to the mitochondrial trifunctional protein (MTP)^42^. This implies that cellular spermidine levels may run low if this pathway is repeatedly triggered such as in heteroplasmy. Spermidine may need to be replenished or else an ‘exhausted type’ of autophagic flux occurs where LC3-II is no longer degraded, as we found here. We also observed that in wild-type cells spermidine inhibits OC differentiation. This is in line with a study that showed supplementation with spermine or spermidine enhances bone strength by inhibiting OCs, whereas inhibiting polyamine biosynthesis reduces these benefits^44^. The differentiation of OC, like that of most cell types, requires autophagy and spermidine may help under homeostatic conditions. In contrast, in heteroplasmic mice, we found that spermidine enhanced OC differentiation at the dose that was previously reported to inhibit the process in wild-type cells^44^. Similar results have been observed in Nrf2 knockout cells^59^, suggesting that spermidine levels may be critically low and need to be replenished under certain stressful situations such as excessive ROS, or repeated clearance of either mitochondria or unwanted complexes I in the cytoplasm.

Our findings demonstrate that spermidine treatment can restore the assembly process and protein import leading to improved mitochondrial function in human *in vitro* OC differentiation, as well as *in vivo* in mice. The m.3243A>G mutation in MELAS patients, which affects a mitochondrial tRNA gene, and the heteroplasmic BL/6^C57-NZB^ mice both exhibit mitochondrial genetic variability, making them comparable models for studying the effects of heteroplasmy. However, it is unclear why the m.3243A>G mutation is pathogenic and why some individuals carrying it are asymptomatic. The human mitochondrial genome is under heavy evolutionary constraint to adapt to the nuclear genome, even without pathogenic mutations^60,61^. A certain threshold of mtDNA mutational load may need to be reached before nucleo-mitochondrial incompatibility arises and MELAS symptoms develop. However, prior work in both mice^8^ and humans^60^ suggests that selection does not always favor nuclear-matching mtDNA variants, indicating forces beyond mitonuclear mismatch. It remains unclear to what extent mitochondrial diseases such as MELAS arise from the accumulation of pathogenic variants leading to mitonuclear mismatch, versus being driven by heteroplasmy itself. Since one of the mouse models in our work features complete mitonuclear mismatch (BL/6^NZB^), we have effectively isolated the influence of heteroplasmy, a distinction that is difficult to achieve in human in vivo samples.

Comparing these murine and human models of disease allowed us to identify shared mechanisms underlying mitochondria-related abnormalities of osteoclastogenesis and led to the discovery of spermidine as a modulator of this process. This finding opens the door for the potential use of spermidine in human heteroplasmic conditions, such as mitochondrial diseases, a therapeutic possibility not previously reported.

## Materials and methods

### Lead Contact and Materials Availability

Further information and requests for resources and reagents should be directed to and will be fulfilled by the Lead Contacts, Anna Katharina Simon and Ana Victoria Lechuga-Vieco.

### Materials availability

Animal lines generated in this study will be made available on request, but we may require a payment and/or a completed Materials Transfer Agreement if there is potential for commercial application.

### Data and code availability

All data reported in this paper will be shared by the lead contact upon request. Original sequencing data files have been deposited at NCBI-SRA (PRJNA1144799), with a temporary link to access the metadata: (https://dataview.ncbi.nlm.nih.gov/object/PRJNA1144799?reviewer=45rhoehhhq1rkj66o92nrv2pds). Links or accession numbers are also listed in the key resources table. The datasets will be publicly available upon the publication of this manuscript. This paper does report original code. The code for 3D bone shape reconstruction/evaluation and OC/OB quantification for in-vitro staining (Supplementary Table 2) can be found in GitHub repository: https://github.com/jianqingzheng/image_processing. Any additional information required to reanalyze the data presented in this work is available from the from the Lead Contact upon request.

## Experimental model and subject details

### Animals

All animal experiments were done in compliance with the guidelines established by the UK Animal (Scientific Procedures) Act 1986 and the Institutional Animal Care and Use Committee of the Barcelona Science Park and University of Barcelona. In vivo experiments with animals were undertaken under the UK Home Office Project License PPL 30/3388. Unless otherwise specified, all experiments were performed in 8- to 12-week-old mice in a C57BL/6JOlaHsd background and a heteroplasmic mouse strain with the nuclear genome of C57BL/6JOlaHsd but mitochondrial DNA of C57+NZB (referred to as BL/6^C57-NZB^)^10^. Mice were housed in specific pathogen-free facilities at the Kennedy Institute of Rheumatology (NDORMS, University of Oxford) and Park Scientific of Barcelona (PCB) under a 12-hour light/dark cycle and had ad libitum access to water and a standard chow diet (SDS diets 801,151; 75% energy from carbohydrates, 17.5% from protein, and 7.5% from fat).

### Patient samples

Patient blood samples were obtained with the approval of the United Kingdom National Research Ethics Service (South Central-Berkshire, approval number 20/04, IRAS 162181). Written informed consent was obtained from all participants, and all procedures were carried out in accordance with relevant national and international guidelines and regulations. This research has been conducted using the UK Biobank Resource. This work was carried out under UK Biobank project number 103356.

## Methods details

### Micro computed tomography (μCT) scan and analysis of bone structure

The PerkinElmer QuantumFX scanner was utilized to scan the tibiae. The region of interest was specified as 100 slices below the growth plate of the proximal tibia and scanned for 3 minutes with an energy of 90KV, 200uA, and field of view of 10. The resulting bone data from the scan were subjected to analysis using Analyse-12.0 software and Analyse-14.0 software.

### Compression testing of femur

After being harvested, the femurs were stored directly in -20 °C. To prepare for the compression test, the femurs were rehydrated with PBS at 4 °C overnight. Murine right femora were subjected to compression test using Electroforce 5500 (TA Instruments, Germany) on a 200 N load. Bones were secured on the lower plate and compressed. For testing, the bones were pre-loaded with a contact force of -0.5 N and load-displacement curves were generated in displacement control at a cross-head speed of 0.3 mm/s until failure of the diaphyseal region. Ultimate load was recorded for each group as an indicator of bone strength.

### Serum and BM tissue processing

On the day of sacrifice, intracardiac puncture was used to collect mouse blood samples using capillary blood collection tubes (16.440, Microvette® CB 300 Z). The collected blood samples were kept on ice for up to 2 hours and then centrifuged at full speed for 5 minutes to extract the serum (supernatant). Serum was stored at −80 °C until further analysis. To collect mouse BM cells, the femurs were crushed with a mortar and pestle in PBS-0.1% BSA-2 mM EDTA. The mixture was then filtered through 70 µm cell strainers, and the collected cells were stored on ice, ready for flow cytometry without RBC lysis.

### ELISA (Enzyme-Linked Immunosorbent Assay)

Mouse serum CTX1 level, mouse serum P1NP level and human serum CTX1 level were measured by ELISA according to the manufacturer’s instructions.

### Tartrate-Resistant Acid Phosphatase (TRAP) Staining

TRAP staining and counterstained with 0.2% methyl green solution experiments were previously described^20^ and conducted according to the manufacturer’s instructions. TRAP positive multi-nuclear (>3 nuclei) OC were counted. The Zeiss AxioScan Z1 Slide Scanner at the Wolfson Imaging Centre, WIMM, University of Oxford was used to acquire the images of OC.

### TRAP staining and histomorphometry analysis of tibiae

Tibiae were fixed in 4% PFA at 4 °C overnight and then decalcified in 10% Ethylenediaminetetraacetic acid (EDTA) decalcification solution for a duration of 14 days (buffer changed every 3 days). After decalcification, the tibiae were paraffin embedded for sectioning using a Leica RM2235 microtome, resulting in sections of 5 µm thickness to create histology slides (VWR 631-1553). The sections were processed and stained with TRAP solution, followed by counterstaining with 0.2% methyl green solution. Histology slides were scanned using the Hamamatsu NanoZoomer S210 (WIMM, the University of Oxford). OC parameters from histology slides were manually quantified by a researcher from JingLianWen Technology (https://www.jinglianwen.com) blinded to the mouse experiment groups.

### Bone Marrow Transplant and chimera blood lineages analysis

To establish BM chimeras, 8-12-week-old female BL/6^C57-NZB^ mice and CD45.1^+^ B6.SJL female mice were lethally irradiated at the dose of 11 Gy (550 cGy twice, 4 hours apart) and rested for 1 hour prior to injection. 1.5 million BM cells derived from 8-12-week-old BL/6^C57-NZB^ and BL/6^C57^ female mice or their respective littermates were transplanted (i.v. injection) into these irradiated mice. Reconstitution effectiveness was evaluated by analysing the percentage of CD45.1^+^, CD45.2^+^ and CD45.1/2^+^ cells in the blood PBMCs of bone marrow-transplanted mice at 2,4, 6- and 8-weeks post-irradiation. To achieve this, we collected 50 microliters of blood from tail vein bleeding and treated it with RBC Lysis Buffer for 5 minutes. After centrifugation, the supernatant was removed, and the cells were stained with antibodies for flow cytometry analysis. Given that previous research^62^ demonstrated that three weeks is sufficient for OC replenishment after transplantation, mice were sacrificed 8-9 weeks following lethal irradiation and transplantation.

This ensured comprehensive reconstitution of most cell lineages with the transplanted cells. Both blood and bone samples were collected for further analysis.

### Mouse In vitro OC differentiation

Mouse in vitro OC differentiation method was previously described^20^. In brief, 8-week-old mouse BM cells were treated with 50 ng/ml M-CSF in Alpha-MEM culture media with 10% FBS, 1% pen/strep, and 1% L-glutamine for 48 hours (day -2) in 96-well plates. On day 0, the adherent cells were differentiated into OCs with 10 ng/ml mouse M-CSF and 50 ng/ml mouse RANKL. Spermidine was added at the dosage of 10µM. Culture media was refreshed every 3 days and OCs were identified by TRAP staining on day 5. The Olympus BX51 microscope at the Kennedy Institute of Rheumatology and AID ELISpot Reader System (ELR078IFL, AID GmbH) at the Wellcome Centre for Human Genetics were used to acquire the images of OC.

### Flow Cytometry

Bone marrow and blood samples were collected from mice and prepared for flow cytometry. Erythrocytes were lysed using eBioscience^TM^ 1x RBC Lysis Buffer. Cells were first incubated with FcR blocker in PBS for 20 minutes, followed by incubation with surface-conjugated antibodies for 30 minutes at 4°C. Conjugated antibodies used for surface staining were: anti-CD45.1, anti-CD45.2, anti-CD45R, anti-CD3e, anti-CD11b, anti-CD117 and anti-CD115. Identification of viable cells was done by fixable Zombie Aqua Live/Dead or near-IR dead cell staining. Stained samples were fixed with BD Cytofix^TM^ Fixation Buffer for 15 minutes at RT. All samples were washed and stored in PBS containing 5 mM EDTA (Sigma-Aldrich) and 2% FBS (Sigma-Aldrich) before acquisition. To determine the absolute cell count within tissues, 10,000 or 20,000 CountBright^TM^ absolute counting beads were added to each sample before flow cytometry to determine the total cell count per bone or blood volume. To ensure accuracy, over 2,000 beads were acquired per tube. Samples were acquired on a FACS LSR II (R/B/V) or Fortessa X20 (R/B/V/YG) flow cytometer (BD Biosciences) using FACSDiva software and analysed with FlowJo (version 10.10, BD Biosciences).

#### Mitochondrial content and ROS levels during OC differentiation

Mitochondrial content and ROS levels were assessed using Nonyl acridine orange (NAO) and CellROX™ Deep Red dyes. Cells were collected and incubated with 100 nM NAO and 5 μM CellROX™ for 20 minutes at 37°C. Ex/Em: 450/530 nm for NAO and 644/665 nm for CellROX™. After washing, cells were surface-stained and prepared for flow cytometry analysis as described in the previous section.

#### Autophagy flux (Flow cytometry)

Autophagy levels were assessed as previously described^39^. In brief, cells were treated for 2 hours with either bafilomycin A1 (10 nM BafA1) or vehicle control, followed by staining using the FlowCellect^TM^ Autophagy LC3 Antibody-based assay kit. After labeling with surface markers (as described earlier), cells were washed with Assay Buffer in 96-well U-bottom plates. Permeabilization was performed with 0.05% saponin, followed by immediate centrifugation. Cells were then incubated with anti-LC3 (FITC) at 1:20 dilution in Assay Buffer for 30 minutes at 4°C, followed by a final wash with 150 µL Assay Buffer. Autophagic flux was determined by calculating the LC3-II median fluorescence intensity as (BafA1 - Vehicle) / Vehicle.

### Pre-processing of single-cell RNA-seq data

To collect mouse BM cells, the femurs were crushed with a mortar and pestle in PBS-0.1% BSA-2 mM EDTA. The mixture was then filtered through 70 µm cell strainers, and the collected cells were plated for in vitro OC differentiation with 50 ng/ml M-CSF in Alpha-MEM culture media. On day 0, the adherent cells were differentiated into OCs with 10 ng/ml mouse M-CSF and 50 ng/ml mouse RANKL. Cells were harvested at the specified time points and were labeled with Cell Multiplexing Oligos (10x Genomics) according to the manufacturer’s instructions. Barcoded sequencing libraries of multiplexed single-cell samples were generated using 10X Genomics Chromium system with 3’ chemistry. Due to a technique issue, two of the three biological replicates were pooled together prior to the multiplexing step and were treated as single replicate (rep2) in the downstream analysis.

### Computational analysis of single-cell RNA-seq data

Raw gene counts were normalized by sequencing depth and log1p transformed in Seurat. Batch effects were corrected using harmony^63^ with n=10,000 variable features, while each sample was treated as an individual batch. Cells were then integrated using n = 30 harmony-corrected principal components, clustered at the resolution of 0.5 and then projected in 2D space using uniform manifold approximation and projection (UMAP). Any clusters that belonged to a non-myeloid lineage or had abnormally high levels of nuclear transcripts (possibly damaged cells with intact nuclei) were discarded, leaving n=10,809 cells for downstream analyses. Cluster marker genes and differentially expressed genes between mouse groups were identified by the Wilcoxon rank-sum test implemented in the Seurat package, and p-values were adjusted using the Benjamini-Hochberg procedure. Pathway enrichment was performed using XGR^64^. Genes associated with oxidative phosphorylation (OXPHOS) were obtained from the KEGG database and were converted to mouse symbols using Ensembl genome database. OXPHOS scores were computed using the “AddModuleScore” function in Seurat and normalized against a set of n=100 random control genes.

The trajectory of osteoclast differentiation was recovered using information from RNA velocity. In brief, genes with a minimum of 20 spliced and 20 unspliced transcripts were extracted from the Cell Ranger alignment outputs and were analyzed in scVelo^22^. The splicing kinetics were recovered using the dynamical model and n=500 top velocity genes. For consistency, we used the same nearest-neighbor graph constructed from scRNA-seq data for all velocity analysis, and inferred velocities were projected onto the scRNA-seq UMAP. The initial and terminal clusters of the differentiation trajectory, together with a latent time for each cell, were inferred using CellRank^23^. We used the default combination of a velocity kernel (80%, deterministic mode) and a connectivity kernel (20%, computed based on the nearest-neighbor graph) to compute the probabilities for cell-cell transitions. The cell-level transitional probabilities were then aggregated at a cluster level and plotted as a fate map using directed PAGA^65^.

For differential abundance test between BL/6^C57^, and BL/6^C57-NZB^ samples, batch-corrected principal components were used to construct single-cell neighborhoods using MiloR^66^, with prop = 0.1, k=20 and d=30. In brief, cells from BL/6^C57^ or BL/6^C57-NZB^ samples were counted within each neighborhood, and the statistical significance of differential abundance was evaluated using a generalized linear model with biological replicates included as covariates. The test results were visualized on a UMAP using the “plotNhoodGraphDA” function.

### Mitochondrial genotyping using single-cell RNA-seq data

Genetic variants in mitochondrial transcripts were recovered at single-cell level using the mgatk pipeline^36^. In brief, sequencing reads were first aligned to a hard-masked mm10 reference genome using CellRanger (v7.1.0; 10x Genomics), followed by mgatk execution to compute the per variant coverage across the mitochondrial genome. The per-cell heteroplasmic ratio was calculated by dividing the summed allelic coverage of all variants specific to NZB mitochondria (n=90, Supplementary Table 1) over the total sequencing coverage at these locations. Only BL/6^C57-NZB^ cells with more than 10 reads covering these variants were retained (n=3,093) for downstream analyses.

### Seahorse analyses

Metabolic assays were performed using a Seahorse XFe96 Analyzer with Seahorse XF Cell Mito Stress Test Kit according to the manufacturer’s instructions. Day 0, Day 1 and Day 3 pre-OCs were seeded in a 96-well plate at the density of 25,000, 25,000and 15,000 cells per well. Cells were treated with 10 ng/ml mouse M-CSF and 50 ng/ml mouse RANKL for indicated time prior to the assay. A final concentration of 1 µM oligomycin, 2 µM FCCP and 1 µM Rotenone and Antimycin A was used based on a pilot titration experiment. Cell density was quantified by CyQuant staining post Seahorse for data normalization.

### ATP Synthesis

ATP levels were quantified using the ATP Bioluminescence Assay Kit CLS II following the manufacturer’s protocol. Briefly, equal numbers of cells from each condition were pelleted, boiled in Tris-EDTA buffer (pH 7.74) for 2 minutes, centrifuged, and the supernatant was analyzed. Luminescence was measured in a CLARIOstar plate reader (BMG LABTECH).

### Analysis of Complex I and Complex II Respiratory Activities

Enzymatic activities were measured spectrophotometrically using the Beer-Lambert law as previously described^67^. Briefly, assays were conducted at 32°C with frozen-thawed mitochondria in activity buffers. Complex I activity was initiated with 130 μM NADH and monitored at 340 nm, subtracting rotenone-insensitive rates. NADH:O₂ and NADH:decylubiquinone activities were assessed in sucrose buffer with specific inhibitors. Complex II activity was measured at 600 nm after adding 10 mM succinate.

### Western Blot Analyses

Autophagy flux during OC differentiation was assessed by incubating the cells in culture media supplemented with 10 nM Bafilomycin A1 for 2 h, with DMSO served as the vehicle control. Autophagic flux was calculated as the ratio of LC3-I to LC3-II.

Cell lysis was performed on ice using NP-40 lysis buffer containing proteinase inhibitors and Phosphatase Inhibitor Cocktail 2. Afterward, the protein concentration was determined using the Pierce™ BCA Protein Assay Kits. The lysate was then treated with Reducing Laemmli Sample Buffer and 15–30 μg of proteins were loaded onto SDS-PAGE gels. For the separation of LC3-I and LC3-II, a 15% Tris-HCl gel and SDS running buffer were employed, while NuPAGE Novex 4– 12% Bis-Tris gradient gels with NuPAGE™ MES SDS Running Buffer (1X) were used for other proteins. Proteins were subsequently transferred to a PVDF membrane and blocked with 5% skimmed milk-PBST. Primary antibodies were incubated overnight in 1% milk, followed by secondary antibody incubation in 1% milk with 0.01% SDS for 1 hour. The following primary antibodies were used: anti-LC3, anti-p62, anti-βActin and Total OXPHOS Rodent WB Antibody Cocktail. Imaging was performed using the Odyssey CLx Imaging System, capturing the signals from IRDye 800CW Donkey Anti-Rabbit IgG (H+L) (1:10,000 dilution) and IRDye 680RD Donkey Anti-Mouse IgG (H+L) (1:10,000 dilution) simultaneously, with varying intensities based on fluorescent exposure. See the Key resources table for a list of primary and secondary antibodies. Data analysis was carried out using Image Studio Lite.

### Confocal imaging

Bone marrow cells were seeded with 10 ng/ml mouse M-CSF and 50 ng/ml mouse RANKL and differentiated for 1, 3, and 5 days in µ-Slide 18-well Ibidi chamber slides (81816, Ibidi GmbH). Cells were fixed with 4% paraformaldehyde for 10 mins and permeabilized with 0.2% Triton X-100 in PBS for another 10 minutes in room temperature. Cells were blocked with PBS containing 2% BSA and 0.01% Tween20 for 1 h and incubated with the primary antibodies overnight at 4 °C. The following antibodies were used to perform immunofluorescence staining in murine and human cells: mouse anti-NDUFA9 (20C11B11B11), anti-NDUFB7 (Polyclonal), anti-ATPβ (3D5), anti-TOMM20 (EPR15581-54). After washing with PBS for three times, cells were incubated in 2% BSA-PBS for 1 hour with the following secondary antibodies: anti-mouse IgG AF488, anti-mouse AF647, anti-mouse AF546, anti-rabbit AF488 and anti-rabbit AF568. After washing with PBS and staining with 2 µg/ml DAPI for 10 minutes, coverslips were mounted in SlowFade™ Diamond Antifade Mountant. Forty to eighty Z-stacks (0.13 µm) were acquired using a Zeiss LSM 980 Airyscan2 confocal system and the ZenBlue software. Data were analyzed using ImageJ/Fiji^68^.

### Genotyping and Heteroplasmy Analysis

Heteroplasmy levels were assessed as previously described^8^. Briefly, mice were genotyped at postnatal day 21 using DNA extracted from ear notches with the REDExtract-N-Amp™ Tissue PCR Kit. Additionally, DNA from cells undergoing OC differentiation (Day 0, Day 1, Day 3, and Day 5) was isolated using 50 mM NaOH at 95°C, followed by neutralization with Tris-HCl (pH 7.5). The G4276A polymorphism, which introduces a BamHI restriction site in C57 mtDNA but not in NZB mtDNA, was used for genotyping. DNA was PCR-amplified with REDExtract-N-Amp™ PCR ReadyMix using primers: Forward 5′-AAGCTATCGGGCCCATACCCCG-3′ (3862-3884) and Reverse 5′-GTTGAGTAGAGTGAGGGATGGG-3′ (4503-4525), under the following conditions: 95°C (30s), 58°C (30s), 72°C (45s) for 30 cycles. The PCR product was digested with FastDigest BamHI at 37°C for 20 minutes and analysed via agarose gel electrophoresis. DNA bands were visualised using the Gel Doc XR+ System (Bio-Rad) and quantified with Quantity One Software. The NZB mtDNA proportion was determined by the ratio of undigested 664-bp fragments to the total of 664-bp (undigested) and 414/250-bp (digested) fragments, with heteroduplex formation corrected using a standard curve from mixed C57 and NZB mtDNA samples.

### Mitochondrial DNA copy number

Total DNA from cells undergoing osteoclast differentiation (Day 0, Day 1, Day 3, and Day 5) was isolated using 50 mM NaOH at 95°C, followed by neutralization with Tris-HCl (pH 7.5). qRT-PCR was performed on Applied Biosystems ®ViiA ™7 Real-Time PCR System using Power SYBR™ Green Master Mix in a 96-well plate to quantify mtDNA copy number relative to nDNA. The primers used were mtCo2 (mtDNA): COII RTF (3’-CTACAAGACGCCACAT-5’, 7037-7052) and COII RTR (3’-GAGGGGGAGAGCAAT-5’, 7253-7238), and SDH (nDNA): SDH RTF (3’-TACTACAGCCCCAAGTCT-5’, 1026-1043) and SDH RTR (3’-TGGACCCATCTTCTATGC-5’, 1219-1202). PCR conditions included an initial denaturation at 95°C for 2 min, followed by 40 cycles of 95°C for 15 s and 60°C for 60 s, with a dissociation curve to confirm specificity. MtDNA copy number was calculated as 2 × (2^ΔCt), with ΔCt = nCt - mtCt, which is inversely proportional to mtDNA content^69^.

### Human osteoclast differentiation

Human peripheral blood mononuclear cells (PBMCs) were isolated from EDTA-blood of healthy donors and MELAS patients. The PBMCs were centrifuged and resuspended in 10% DMSO in FBS and stored in -80° C. On the day of seeding (day -3), 2.5 × 10^5^ cells/well of human PBMCs were seeded in 96-well plates with 100 μL/well of Alpha-MEM culture media containing 100 ng/ml human M-CSF, 10% FBS, 1% pen/strep, and 1% L-glutamine. This setup was maintained at 37°C in a 5.5% CO2 humidified incubator for 72 hours to facilitate the purification of adherent cells through plastic adhesion. On day 0, the cells were induced to differentiate into OCs in 100 μL/well of complete osteoclastogenic media (Alpha-MEM culture media with 10% FBS, 1% pen/strep, and 1% L-glutamine) supplemented with 50 ng/ml human M-CSF and 100 ng/ml human RANKL, and spermidine at 10µM. Subsequently, the culture medium was refreshed on day 2 and day 4. By day 6, mature OCs had formed and were fixed using 4% PFA for TRAP analysis.

### Statistical analysis

The mean ± standard error of the mean (SEM) was used to represent all data. When comparing two experimental groups, a two-tailed t-test was employed. For datasets that were normally distributed with equal variances, an unpaired two-tailed Student’s t-test was utilized. In experiments that had ≥3 experimental groups, one-way analysis of variance (ANOVA) and multiple comparisons with Tukey’s correction were employed. Paired or unpaired one-way ANOVA was utilized for multiple comparisons of normally distributed datasets with one variable. The p-value was applied to evaluate the statistical significance of the hypothesis being tested. GraphPad Prism (San Diego, CA, Version 9.5.1 (528)) was used for statistical analyses. Wilcoxon rank-sum tests were used in single-cell statistical testing in R (v4.2.2).

### 3D bone reconstruction

To reconstruct and visualize the bone, Algorithm 1 is used (Supplementary Table 1). First, the CT volumes are rotated and translated to a standard position and orientation. Next, binary volumes of the entire skeleton structure and the cortical bones are extracted from the input volumes through thresholding. Subsequently, the connected components in the binary volumes are counted. This step aids in removing the fibular bone and retaining the largest connected cortical bone, as facilitated by Algorithm 2. Thereafter, the largest cortical bone is dilated and then eroded to fill the hole inside the tibia using Algorithm 3. In the final step, the cortical bones with filled holes intersect with the tibial skeleton structure, excluding the fibula, to extract the 3D shape of the tibial bone in the region of interest.

In Algorithm 2, a binary volume *V^i^* and the index *k* serve as inputs (Supplementary Table 2). The initial step involves identifying and counting the connected components using MATLAB’s built-in function “bwconncomp()”. In the subsequent step, the connected components within the binary volume are labeled and tallied using MATLAB’s built-in functions “labelmatrix()” and “regionprops()”. These connected components are then arranged in descending order using MATLAB’s “sort()” function. The value of index *k* is assessed to determine whether it’s positive or negative. If *k* is positive, the *k^t^*^ℎ^ largest component is retained while others are discarded; if *k* is negative, the *k^t^*^ℎ^ largest component is eliminated. The algorithm concludes by outputting the remaining components, *V°*. As a result, this methodology is adept at removing the fibular bone and any extraneous fragments that lie outside the tibial bone.

Algorithm 3 takes a binary volume *V^i^* and a series of iteration numbers for dilation or erosion [*t*_1_, *t*_2_, ⋯ *t_k_*] as inputs (Supplementary Table 2). The structuring element is initialized using MATLAB’s built-in function “strel()”. Subsequently, for each iteration number designated for dilation or erosion, the iteration number *t* is examined. If *t* is positive, the copied binary volume *V°* undergoes *t* dilation iterations, and if *t* is negative, *V°* is eroded for *t* times. This algorithm is thus integrated into Algorithm 1 with the aim of filling holes inside the cortical bones and subsequently compressing the volume to adhere to the original bone size.

### Quantification of OC from in vitro staining

To quantify the OC parameters from microscopic images, Algorithm 4 is employed to determine the area size and the ratio of the cell region. In Algorithm 4, each image is processed using its corresponding mask for the region of interest and specific thresholding values (Supplementary Table 2). The algorithm starts by normalizing the intensity of the input images. Subsequently, the red channel of the image is extracted through thresholding to identify and isolate the cell areas. The next step involves identifying and counting the connected components with MATLAB’s built-in function “bwconncomp()”. Following that, the connected components within the binary image are labeled and enumerated using MATLAB’s “labelmatrix()” and “regionprops()” functions. For each connected component, excluding the background region, the area size and its ratio in the entire region of interest are computed based on the areas of the red channels.

## Acknowledgements

We acknowledge all Autophagy in the Immune System group members for their scientific discussions contributing to this manuscript and Prof. Jose Antonio Enríquez (Spanish National Center for Cardiovascular Research, CNIC, Madrid) who provided mouse models for our research and providing thoughtful feedback. We thank Dr. Hanlin Zhang from University of California, Berkeley for discussions and constructive comments. We thank Jianwei Cui from the University of Oxford who helped with experiments. We thank Patricia Cotta Moreira and Mino Medghalchi from the Kennedy Institute of Rheumatology animal facility, and the Barcelona Science Park (PCB) animal facility for their excellent care and assistance of animal well-being. We thank Thomas Conrad, Caroline Braeuning and Sarah Nathalie Vitcetz at the MDC Genomics Platform (BIH, Charité Campus Mitte, Berlin) for discussions about transcriptomic experiments and sequencing performed at their facility. Histology was performed in the Kennedy Institute of Rheumatology Histology Facility by Dr. Ida Parisi. We obtained confocal microscopy assistance from Christoffer Lagerholm, Senior Advanced Microscopy Manager at the Kennedy Institute of Rheumatology, and the Advanced Digital Microscopy (ADM) Core Facility at IRB Barcelona is greatly appreciated. We thank the NHS Highly Specialised Service for Rare Mitochondrial Disorders (Oxford University Hospitals, UK) for patient recruitment and for sharing clinical samples. We thank Dr. Elisabeth Greßler from Max-Delbrück-Center for Molecular Medicine for experimental assistance. Images were created using BioRender.com under the academic license granted to AKS.

This work was supported by: University of Oxford Medical and Life Sciences Translational Fund MC_PC_17174 and MC_PC_18059 from Wellcome ISSF fund to HL. China Scholarship Council fund (202006320024) to JJ. UMDF PF-23-0010 to RJ-M. Biotechnology and Biological Sciences Research Council (BBSRC, BB/X007049/1) to SM. Kennedy Trust Prize Studentship (AZT00050AZ04) to JZ. Chinese Academy of Medical Sciences (CAMS) Innovation Fund for Medical Sciences (CIFMS) 2018-I2M-2-002 to LL. Centre for OA Pathogenesis (Versus Arthritis 20205) to TLV. KTRR KENN202111 to AW. Oxford Hospitals charity and Lily Foundation “Treatments for Mitochondrial Disease – 2019/20” to JP. Salary support from the UK NHS Specialist Commissioners who fund the “Rare Mitochondrial Disorders of Adults and Children” clinical Service to SDW. Novo Nordisk Foundation (NNF0064142) to COC. This work is funded by Diabetes UK (19/0005994 and 21/0006335), MRC (MR/T00200X/1) and Wellcome’s Institutional Strategic Support Fund awarded to University of Exeter; KAP and TH is funded by Wellcome Trust (219606/Z/19/Z); The work is supported by the National Institute for Health Research (NIHR) Exeter Biomedical Research Centre, Exeter, UK. European Molecular Biology Organization Postdoctoral Fellowship (EMBO) ALTF115-2019, “La Caixa” Foundation (ID 100010434, LCF/BQ/PI24/12040005), the Fundación Ramón Areces grant (Grant No. CIVP22A7576), and funding from MINECO through the Centers of Excellence Severo Ochoa Award and CERCA Programme of the Generalitat de Catalunya to AVL-V. Wellcome Trust Fund 220784/Z/20/Z to AKS.

## Author contributions

AKS, A.V.L.-V. and H.L. conceptualized and designed the study.

A.V.L.-V., H.L., and J.J. performed and analyzed experiments, methodology, and investigation.

AKS, A.V.L.-V., H.L. and J.J. discussed analyses.

AKS and A.V.L.-V. supervised the project and interpreted the experimental data.

AKS, A.V.L.-V., H.L. and J.J. wrote the original draft.

AKS, A.V.L.-V., H.L., J.J. and TLV reviewed and edited the manuscript.

C.OC. planned experiments, analyzed data and edited the final manuscript.

K.G. performed mitochondrial DNA copy numbers, respiratory complexes activities and confocal imaging.

R.J-M. performed respiratory complexes analyses.

T.H. and K.A.P. investigated the association between m.3243A>G and bone parameters in individuals from the UK Biobank cohort.

J.Z. and TLV contributed to the 3D reconstructions of bone images and image analysis.

L.L. performed the isolation of donor PBMC samples, followed by OC differentiation, and autophagy flux assays.

S.W. and A.W. analyzed the bone stiffness parameters.

S.D.W. and J.P. provided samples and contributed to discussions on human and patient data. All authors reviewed and approved the final manuscript.

---

**Extended Data Figure 1.**
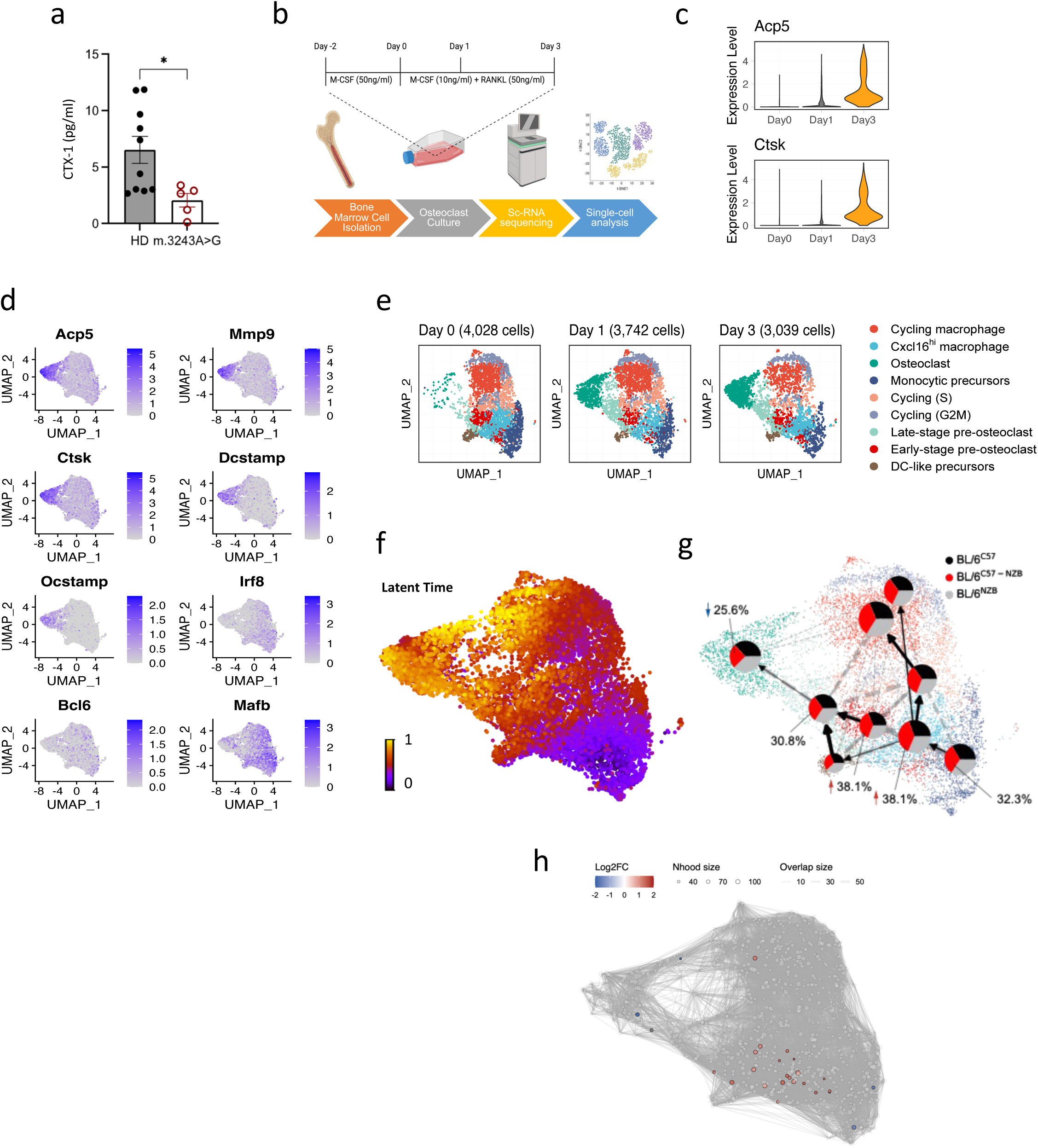
Single-cell transcriptomic profiling and trajectory analysis of OC formation. **a** Quantification of bone resorption marker CTX-1 levels in human serum samples from healthy donors (HD, n=10) and m.3243A>G patients (n=5). *p < 0.05, two-tailed unpaired t-test. Data are presented as mean values ± SEM. **b** Schematic of experimental strategy for single-cell transcriptomics of murine OC formation. Bone marrow cells from BL/6^C57^ (n=3), BL/6^NZB^ (n=3), and BL/6^C57-NZB^ (n=3) strains were isolated and cultured with M-CSF (50 ng/ml) for 2 days, followed by M-CSF (10 ng/ml) and RANKL (50 ng/ml) from day 0 to day 3. Single cell sequencing and analysis were then performed with day 0, day 1 and day 3 cells. **c** Expression levels of OC marker genes (*Acp5* and *Ctsk*) from aggregated single cells at each time point, indicating successful *in vitro* osteoclastogenesis. **d** Uniform manifold approximation and projection (UMAP) visualization of the expression levels of OCs marker genes (*Acp5*, *Mmp9, Ctsk, Dcstamp,* and *Ocstamp*) and negative regulators (*Irf8*, *Bcl6*, and *Mafb*). Color indicates log-normalized gene expression levels. **e** UMAP visualization of single-cell transcriptomic profiles (n = 10,809 cells) during OC differentiation. Cells are clustered by transcriptomic similarity and shown at three time points: day 0, day 1 and day 3. **f** UMAP projection of velocity-latent time estimated from the RNA splicing dynamics. Cells with a latent time of 0 or 1 represent the starting or terminating population inferred by RNA velocity, respectively. **g** Fate map for OC differentiation using directed probabilistic approximate graph abstraction (PAGA). Each node represents one cluster, and arrows represent aggregated velocity flow concordant to the velocity-inferred trajectory. Dotted lines represent other velocity flow between each pair of clusters. Pie charts denote the relative abundance of BL/6^C57^, BL/6^NZB^ and BL/6^C57-NZB^ cells, normalized by the total number of cells in each group. The relative abundance of BL/6^C57-NZB^ cells for each cluster along the OC differentiation trajectory is labelled next to the pie chart. **h** Differential abundance test representation. The relative enrichment of BL/6^C57-NZB^ cells in early-stage pre-OCs (shown in red), as well as the relative depletion in mature OCs (shown in blue), indicates premature termination of osteoclastogenesis in heteroplasmic cells. Nodes represent neighborhoods of cells with similar transcriptomic profiles, colored by the relative abundance (log2FC) of BL/6^C57-NZB^ cells compared to BL/6^C57^. Node sizes correspond to the number of cells in a neighborhood. Non-significant neighborhoods (p>0.05) are colored in white and plotted in the background. The statistical significance of this observation was validated by differential abundance testing using MiloR.

**Extended Data Figure 2.**
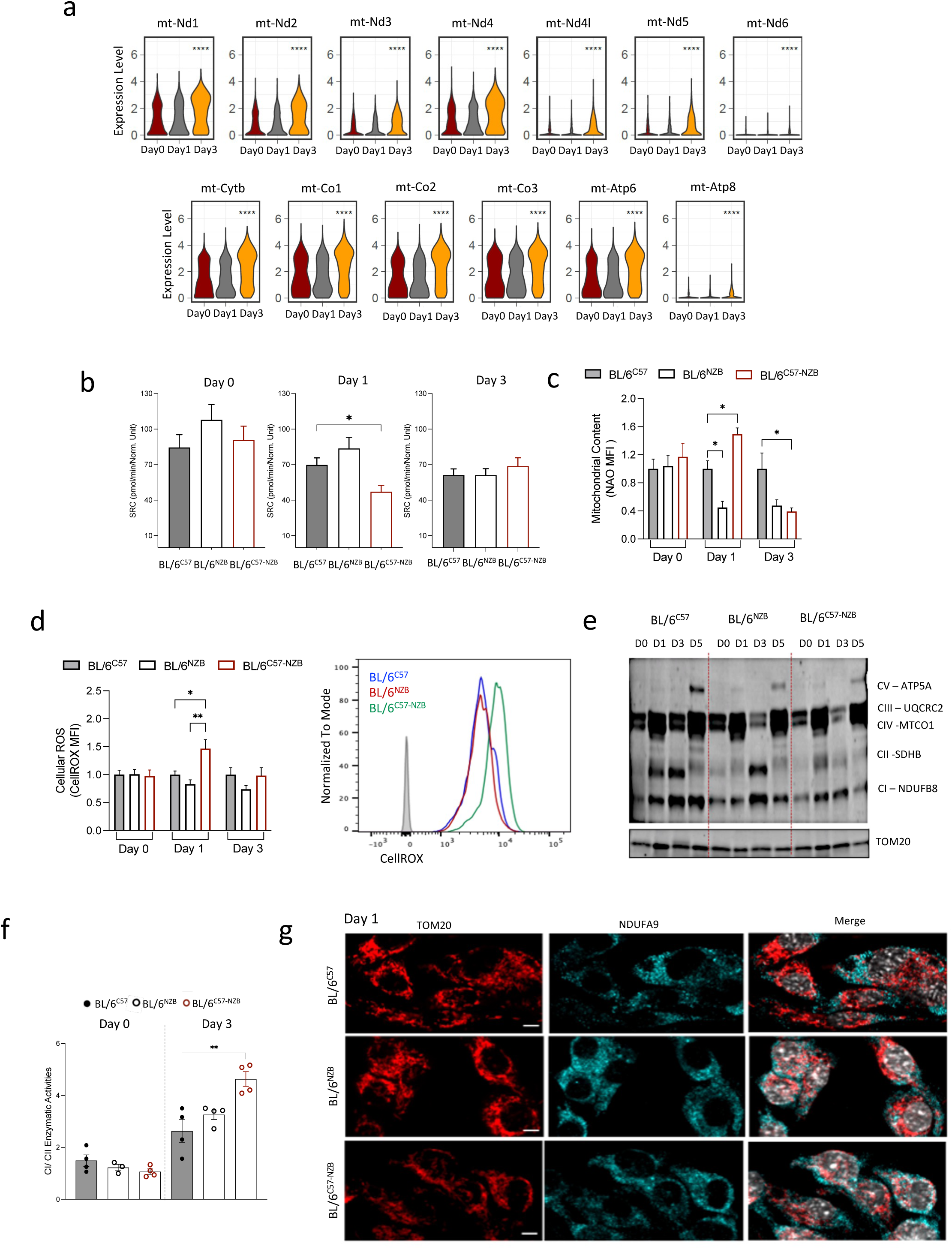
Analysis of mtDNA-encoded genes, respiratory complexes, and bioenergetic parameters during OC differentiation. **a** Violin plots displaying the average expression levels of mitochondrial DNA-encoded genes in BL/6^C57^ cells at day 0 (n = 1,550), day 1 (n = 1,148), and day 3 (n = 1,292) during OC differentiation. Complex I: mt-Nd1, mt-Nd2, mt-Nd3, mt-Nd4, mt-Nd4L, mt-Nd5 and mt-Nd6. Complex III: mt-Cytb. Complex IV: mt-Co1, mt-Co2 and mt-Co3. Complex V (ATPase): mt-Atp6 and mt-Atp8. The y-axis represents the depth-normalized gene expression level in log scale (log count per 10,000 sequencing reads). **** p < 0.0001, Wilcoxon test, with day 0 and day 1 cells as the reference groups. **b** Spare respiratory capacity (SRC) calculated by subtracting basal respiration from maximal respiration during *in vitro* OC differentiation (day 0 to day 3) for the indicated genotypes (n=5, two independent experiments). *p < 0.05, two-tailed Dunnett’s test following one-way ANOVA. Data are presented as mean ± SEM. **c** Mitochondrial content was assessed using N-acridine orange (NAO) staining. Live cells were analyzed by FACS during OC differentiation from day 0 to day 3. n= 3 BL/6^C57^ n=3 BL/6^NZB^ and n=5 BL/6^C57-NZB^. * p<0.05, determined using Tukey’s multiple comparisons following two-way ANOVA. Data are normalized to wild-type BL/6^C57^ and presented as mean ± SEM. **d** ROS quantification on day 0, day 1 and day 3 during OC differentiation. Left: Quantification of CellROX Deep Red MFI measurements. Right: FACS histogram plot are representative measurements on day 1 from cells derived from BL/6^C57^ (blue), BL/6^NZB^ (red) BL/6^C57-NZB^ (green) mice, where the grey line shows background MFI (unstained). n= 3 BL/6^C57^ n=3 BL/6^NZB^ and n=5 BL/6^C57-NZB^. * p<0.05, ** p <0.01 determined using Tukey’s multiple comparisons following two-way ANOVA. Data are normalized to wild-type BL/6^C57^ and presented as mean ± SEM. **e** Western blot analysis for OXPHOS complexes protein expression in day 0, day 1 and day 3 of *in vitro* OC differentiation. CI-NDUFB8, CII-SDHB, CIII-UQCRC2, CIV-MTCO1 and CV-ATP5A levels are shown. TOM20 expression was used as mitochondrial protein loading control. Representative image from four independent experiments, n=4 mice per genotype. **f** Ratio of Complex I to Complex II activities. Respiratory complexes activities were measured spectrophotometrically using 5 µg of total mitochondrial protein per assay. Measurements were performed in four replicates using pre-OCs on days 0 and 3 of differentiation. n= 4 mice per genotype. ** p <0.01, Dunnett’s test following one-way ANOVA. Data are presented as mean ± SEM. **g** Confocal imaging of mitochondrial TOM20 (red), Complex I subunit NDUFA9 (cyan) and DAPI (grey) in pre-OCs on day 1 of differentiation derived from BL/6^C57^, BL/6^NZB^, and BL/6^C57-NZB^ mice. Representative confocal images for the indicated markers. Scale bar: 5 μm, n = 4 mice per genotype.

**Extended Data Figure 3.**
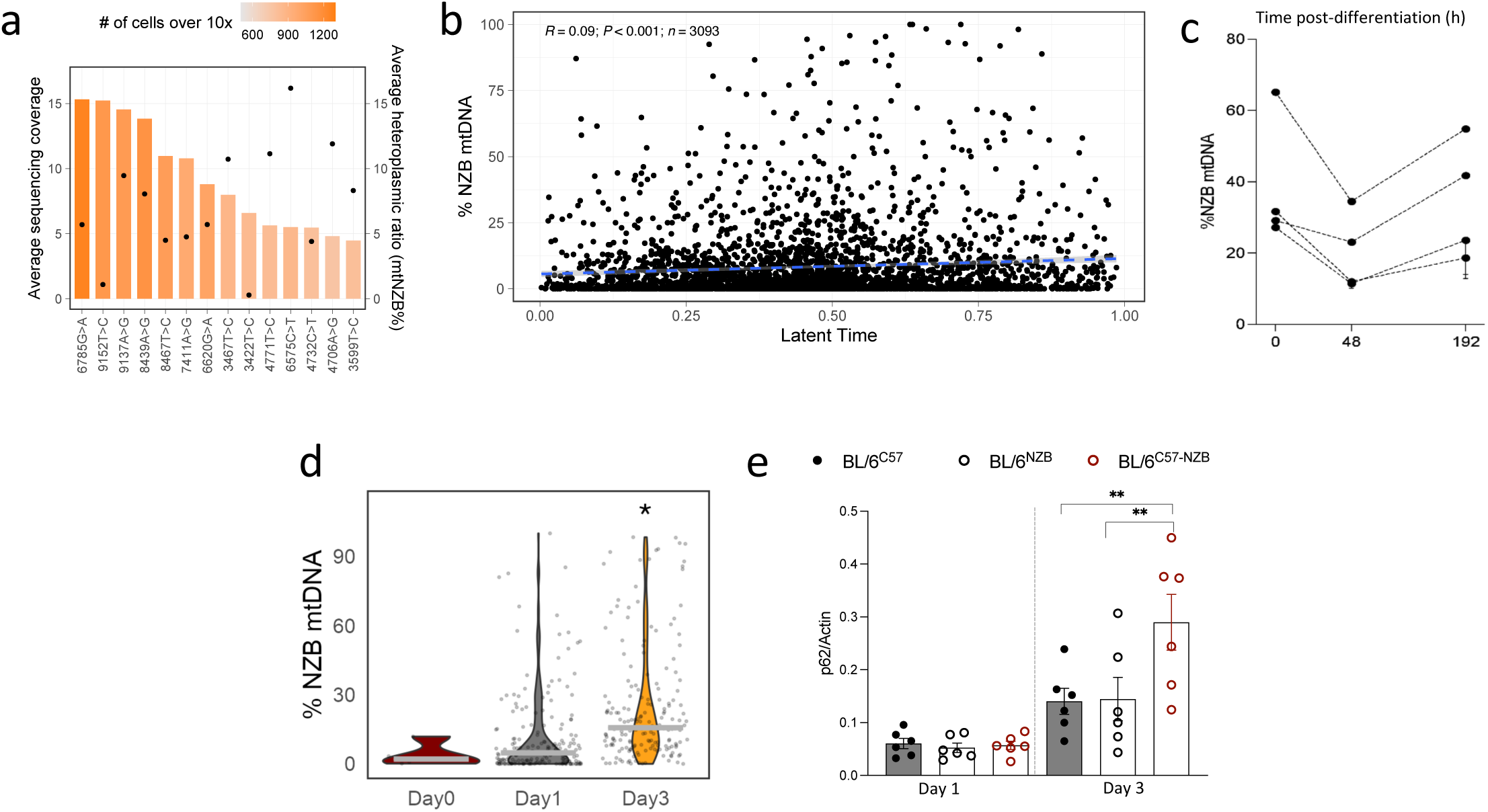
Analysis of mitochondrial selection and the role of autophagy during OC maturation. **a** Average sequencing depth (bars) and the average heteroplasmic percentage (dots) for selected NZB mtDNA variants in heteroplasmic cells. Colors represent the number of cells with at least 10x coverage for a given variant. **b** NZB mtDNA levels (%NZB mtDNA, y axis) plotted against the velocity latent time (x axis, same latent time as shown in **Extended Data Fig.1f**), demonstrating a slight shift towards NZB mtDNA during OC differentiation. *** p < 0.001, Spearman’s rank correlation test. **c** Assessment of the percentage of NZB mtDNA (% NZB mtDNA) during *in vitro* OC differentiation of heteroplasmic cells (0, 48, and 192 hours after the addition of 10 ng/ml M-CSF and 50 ng/ml RANKL). NZB mtDNA levels were quantified by PCR, followed by RFLP (Restriction Fragment Length Polymorphism) to distinguish between C57 and NZB mtDNA variants. Samples were derived from n = 4 BL/6^C57-NZB^ mice. **d** NZB mtDNA levels (%NZB mtDNA, y axis) in the OC cluster (cluster 2) at day 0, day 1 and day 3 of differentiation. * p < 0.05, Wilcoxon Rank Sum test. **e** Quantification of p62 expression from Western blot analyses shown in **Fig.4d**. Actin was used as a protein loading control. n = 6 independent biological samples for DMSO control and BafA1; each dot represents an individual mouse. **p < 0.01, determined using Tukey’s multiple comparisons following two-way ANOVA. Data are presented as mean values ± SEM.

**Extended Data Figure 4.**
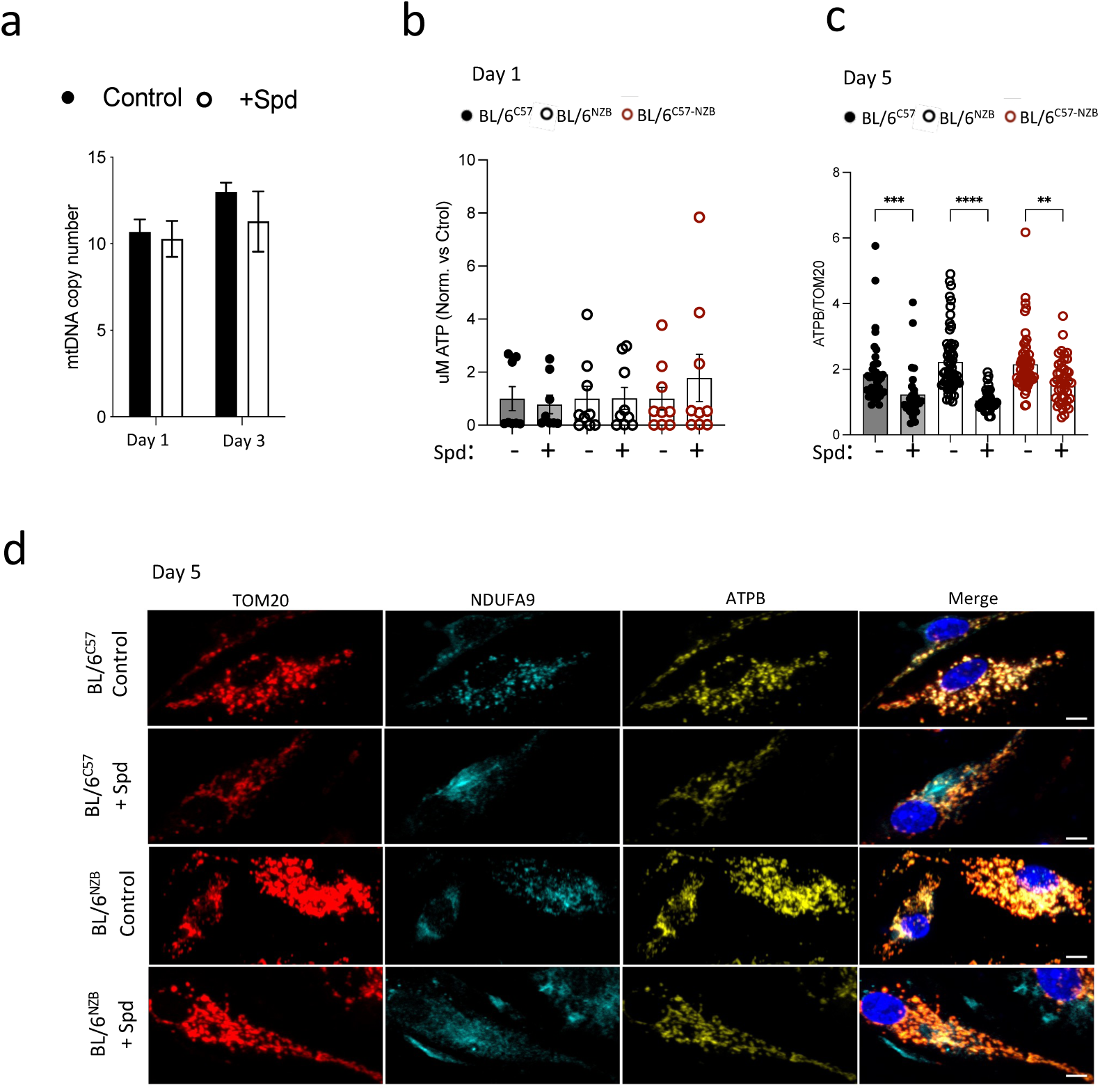
Spermidine enhances mtDNA selection and complex I expression in heteroplasmic cells. **a** Mitochondrial DNA copy numbers in BL/6^C57-NZB^ cell cultures without (Control) or with 5 µM spermidine treatment (+Spd) on day 1 and day 3 during OC differentiation (n=4-5 mice). MtDNA copy numbers were normalized to genomic DNA copy numbers, as determined by qPCR (2−ΔCT). Data are presented as mean values ± SEM. **b** Total ATP levels were measured by luminometry on day 1 of OC differentiation in cells derived from BL/6^C57^, BL/6^NZB^, and BL/6^C57-NZB^ mice, with and without 10 µM spermidine treatment. Data are normalized to untreated cells and each dot represents a sample from an individual mouse of the indicated genotype. Data are presented as mean values ± SEM. **c-d** Confocal imaging of mitochondrial TOM20, Complex I subunit NDUFA9, and ATPase subunit ATPβ in OCs derived from BL/6^C57^and BL/6^NZB^, and BL/6^C57-NZB^ mice with and without spermidine treatment on day 5 of *in vitro* OC differentiation. **c,** Protein quantification ratio of ATPβ per mitochondrion (ATPβ/TOM20), based on marker signals (Integrated Density, IntDen) (n=3 mice per genotype). **** p <0.0001, *** p <0.001, ** p <0.01, two-tailed Dunnett’s test following one-way ANOVA. Data are presented as mean values ± SEM.**d,** Representative confocal images for the indicated markers in OCs derived from BL/6^C57^and BL/6^NZB^ mice with and without spermidine treatment on day 5 of *in vitro* OC differentiation. Scale bar: 5 μm.

**Extended Data Figure 5.**
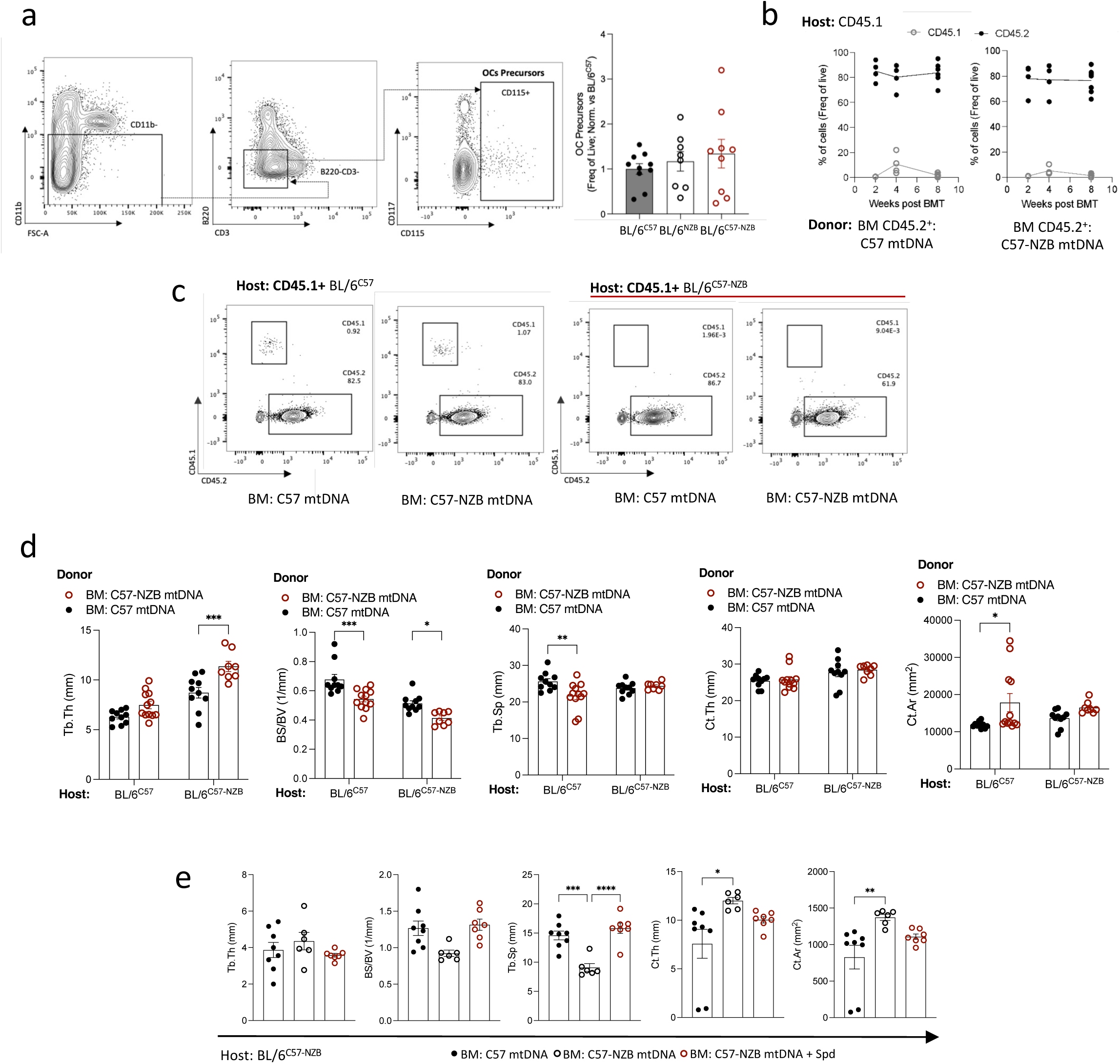
OC progenitor gating, chimera reconstitution, and bone analysis. **a** Flow cytometry gating strategy of the OC progenitors (CD11b-B220-CD3-CD115+) in BM cells (left) and percentage of OC progenitors (right) from BL/6^C57^ mice (n=10), BL/6^NZB^ mice (n=8) and BL/6^C57-NZB^ (n=9) mice. Data are presented as mean ± SEM. **b** Assessment of reconstitution efficacy in chimeric mice by evaluating the BM CD45.2+ population. Each dot represents an individual animal. Filled dots: donor cells; Empty dots: endogenous cells. **c** Representative flow cytometry plots showing the gating strategy and distribution of CD45.1+/CD45.2+ BM cells from the specified groups. **d** Morphological bone parameters in tibiae measured by micro-CT. Tb.Th, BS/BV, Tb.Sp, Ct.Th and Ct.Ar (BL/6^C57^ to CD45.1+ B6.SJL group (n=10); BL/6^C57-NZB^ to CD45.1^+^ B6.SJL group (n=12); BL/6^C57^ to BL/6^C57-NZB^ group (n=10); BL/6^C57-NZB^ to BL/6^C57-NZB^ group (n=8). Each dot represents and individual animal. *p < 0.05, **p < 0.01; ***p < 0.001, determined by two-way ANOVA test. Data are presented as mean ± SEM. **e** Lethally irradiated young BL/6^C57-NZB^ mice were reconstituted with bone marrow from BL/6^C57^ or BL/6^C57-NZB^ mice and treated with 5 mM spermidine in drinking water for 8 weeks. Tb.Th, BS/BV Tb.Sp, Ct.Th and Ct.Ar of tibiae quantification by micro-CT (n = 8 BL/6^C57^; n = 6 BL/6^C57-NZB^ and n = 7 BL/6^C57-NZB^ + Spd; each dot represents an individual animal). The experiment is representative of two independent experiments. *p < 0.05, **p < 0.01, ***p < 0.001, ****p < 0.0001, one-way ANOVA followed by Tukey’s multiple comparisons test.

**Supplementary Table 1:**
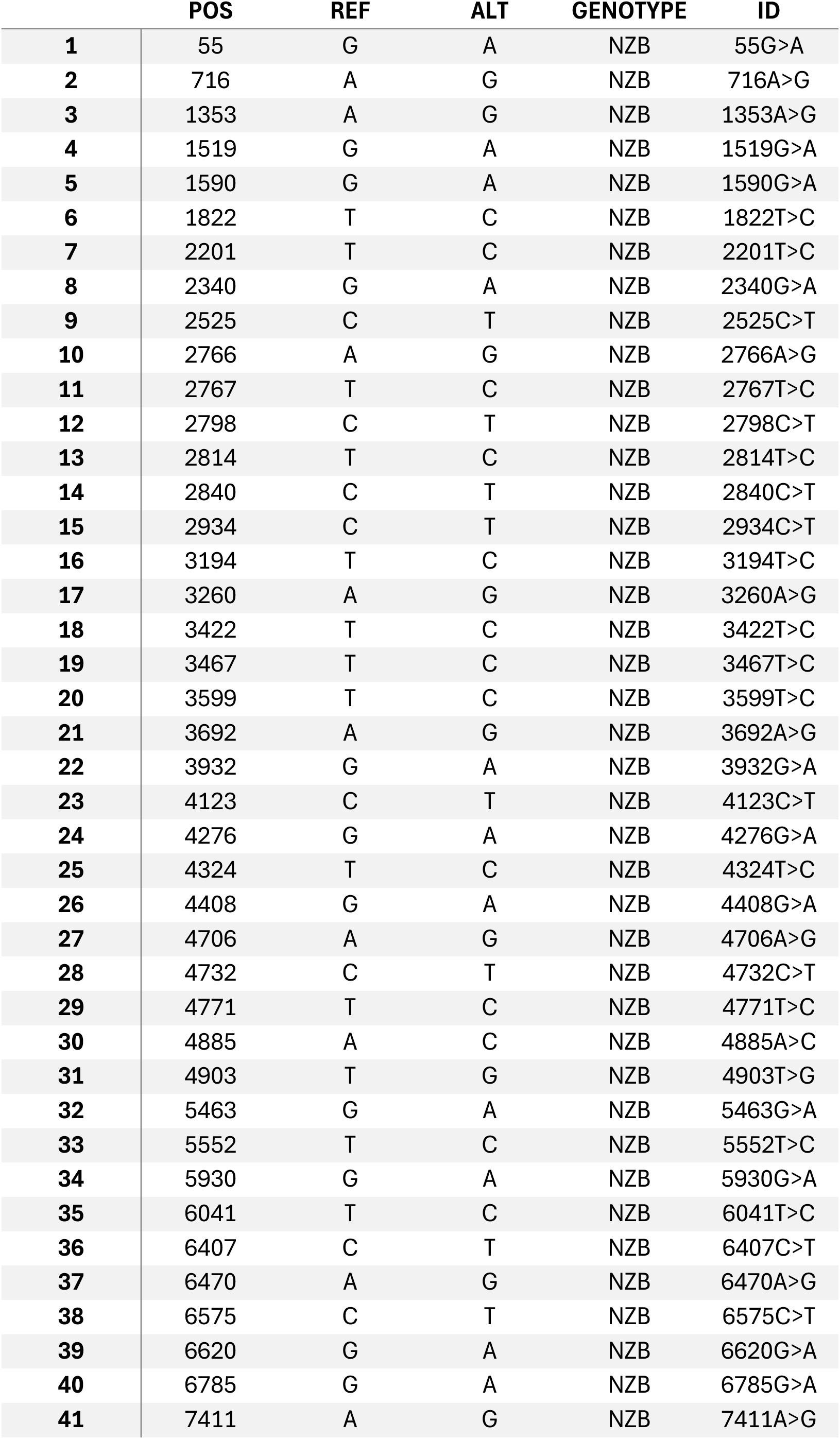
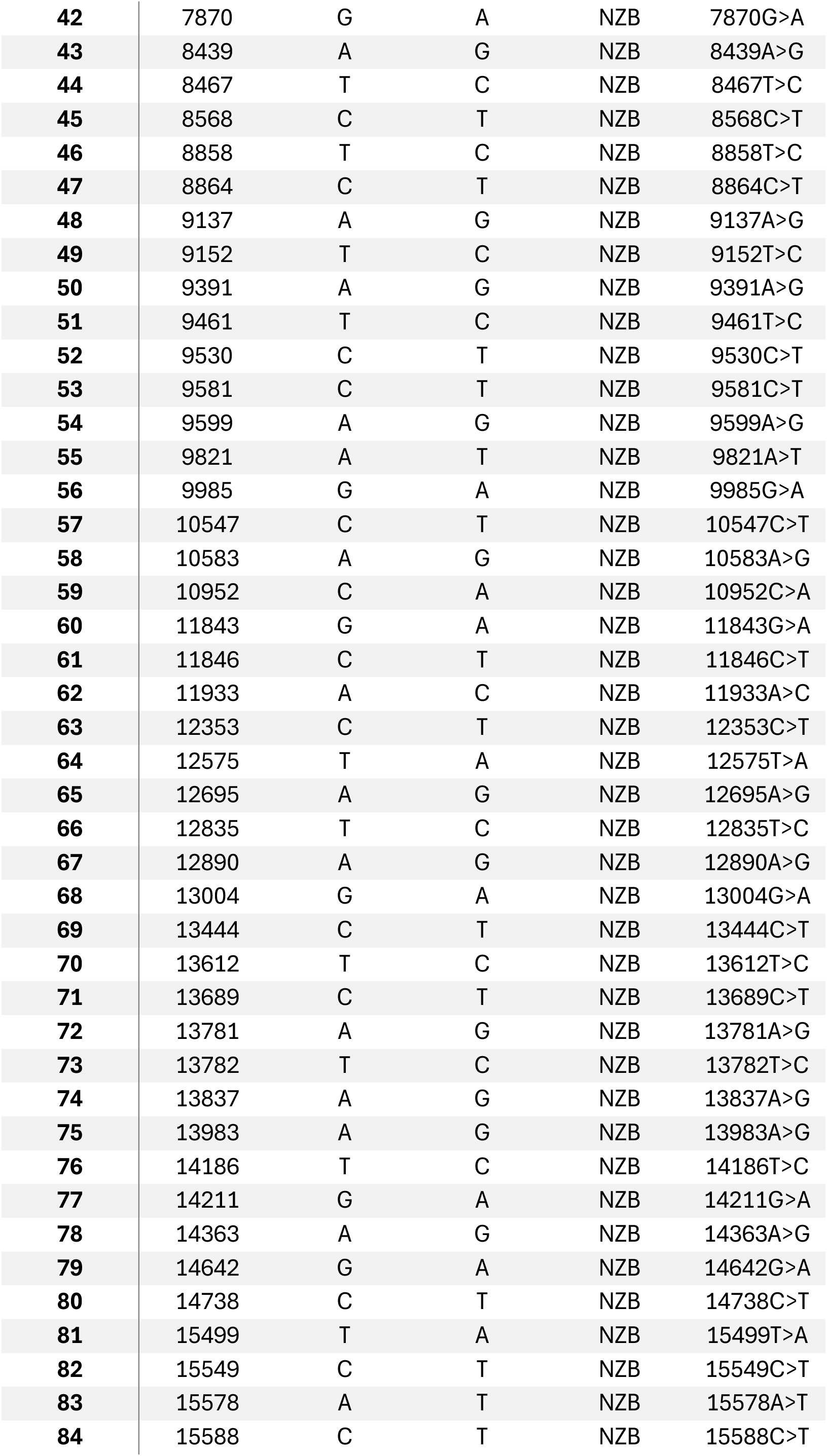
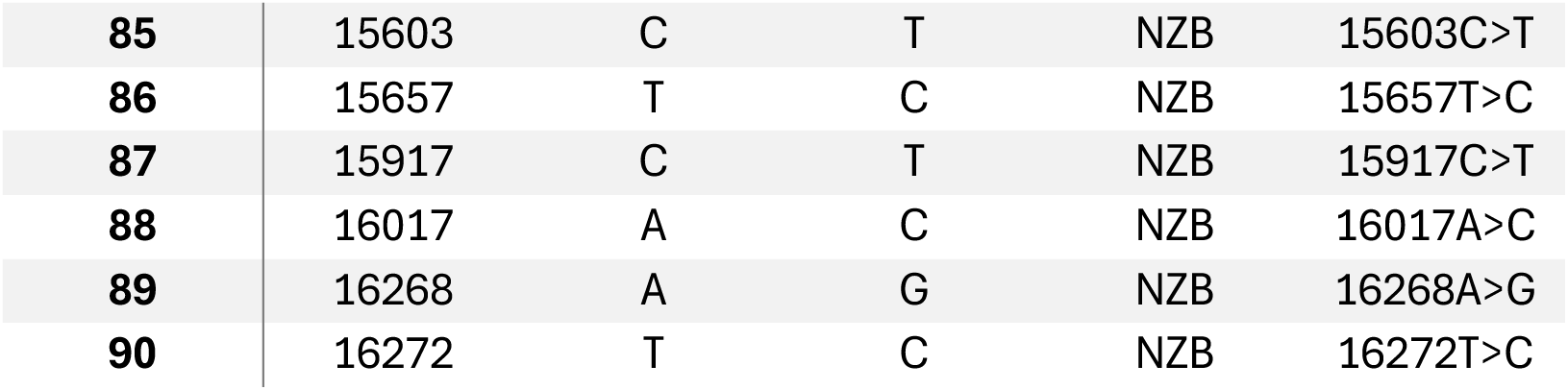
NZB mtDNA SNPs identified from scRNA-seq data. The per-cell heteroplasmic ratio was calculated by dividing the summed allelic coverage of all variants specific to NZB mitochondria (n=90) over the total sequencing coverage at these locations.

**Supplementary Table 2:**
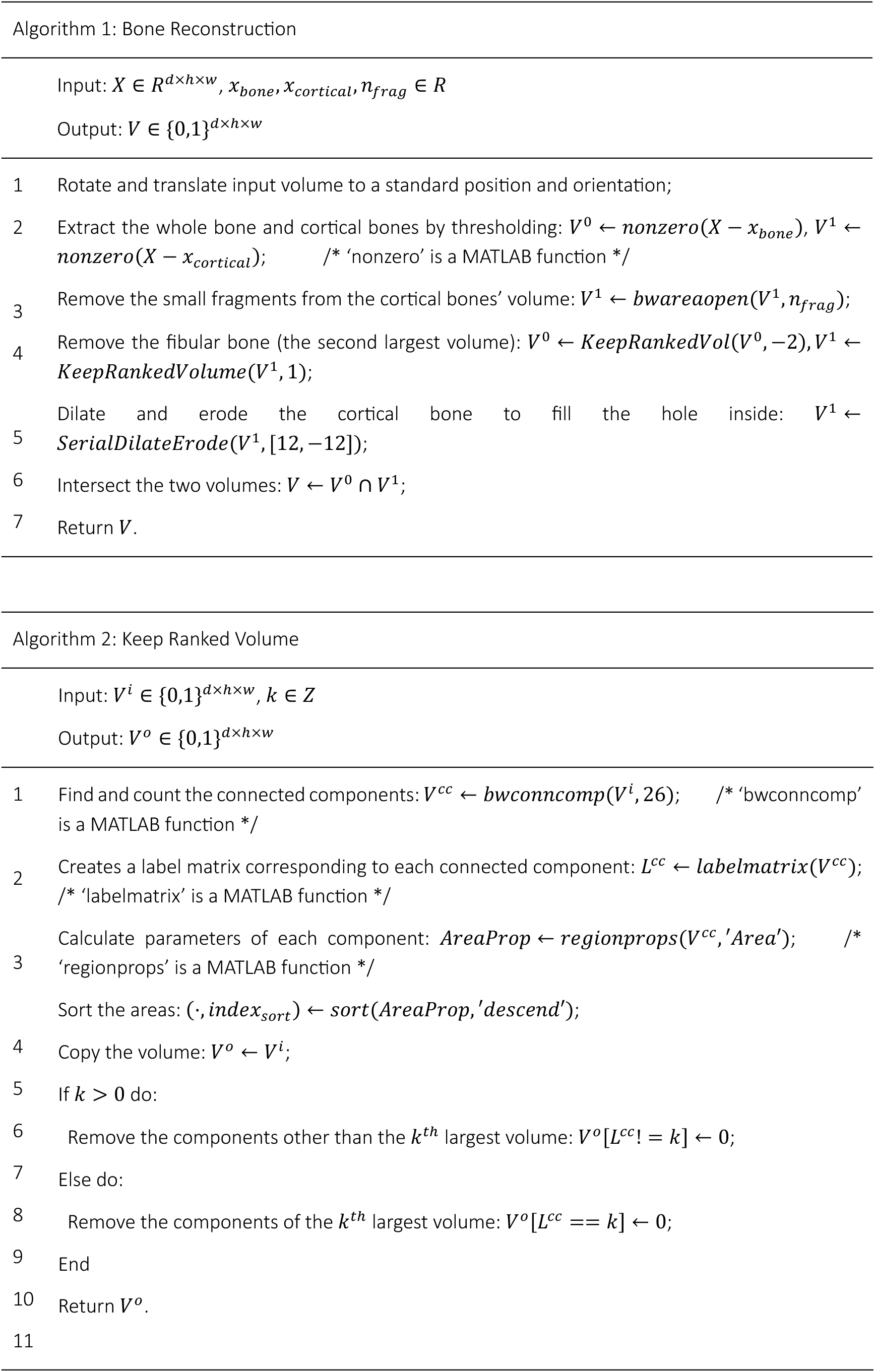
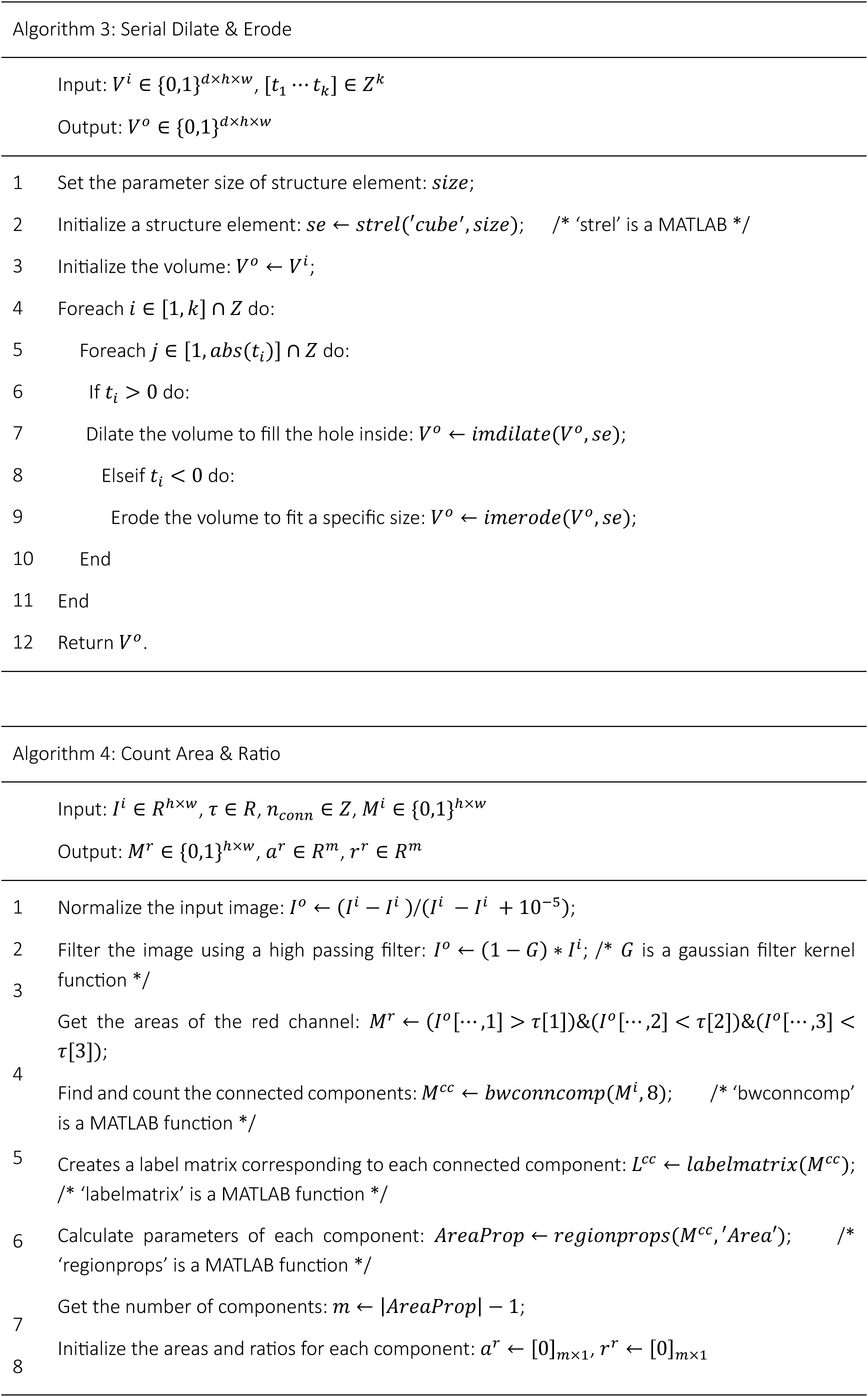
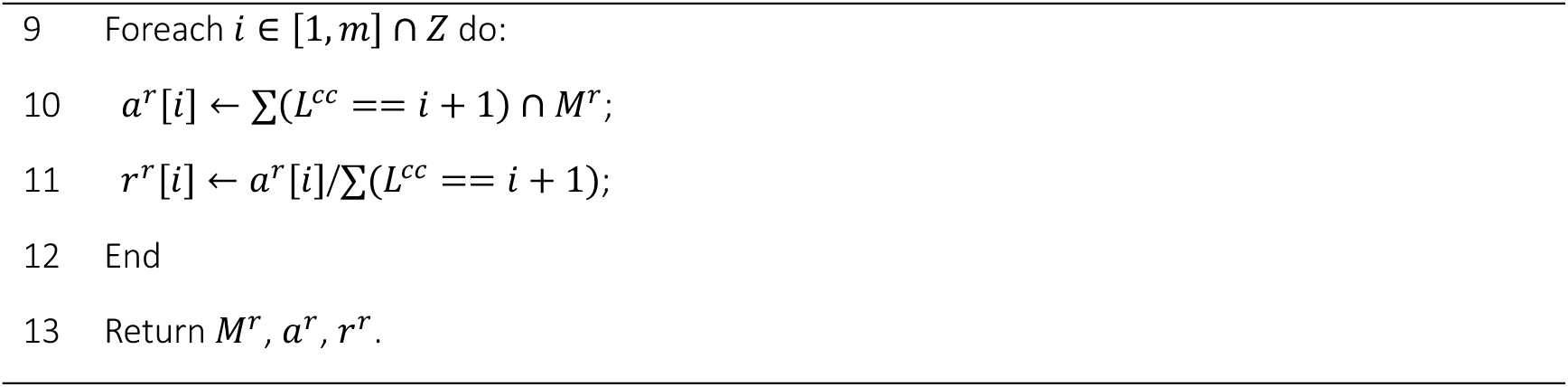
Algorithms used for bone reconstruction, keep ranked volume, Serial Dilate & Erode, Count Area & Ratio.

| Algorithm 1: Bone Reconstruction |  |
|---|---|
|  | Input: X ∈ R<sup>d×h×w</sup>, x<sub>bone</sub>, x<sub>cortical</sub>, n<sub>frag</sub> ∈ R |
|  | Output: V ∈ {0,1}<sup>d×h×w</sup> |
| 1 | Rotate and translate input volume to a standard position and orientation; |
| 2 | Extract the whole bone and cortical bones by thresholding: V<sup>0</sup> ← nonzero(X − x<sub>bone</sub>), V<sup>1</sup> ← nonzero(X − x<sub>cortical</sub>); /* ‘nonzero’ is a MATLAB function */ |
| 3 | Remove the small fragments from the cortical bones’ volume: V<sup>1</sup> ← bwareaopen(V<sup>1</sup>, n<sub>frag</sub>); |
| 4 | Remove the fibular bone (the second largest volume): V<sup>0</sup> ← KeepRankedVol(V<sup>0</sup>, −2), V<sup>1</sup> ← KeepRankedVolume(V<sup>1</sup>, 1); |
| 5 | Dilate and erode the cortical bone to fill the hole inside: V<sup>1</sup> ← SerialDilateErode(V<sup>1</sup>, [12, −12]); |
| 6 | Intersect the two volumes: V ← V<sup>0</sup> ∩ V<sup>1</sup>; |
| 7 | Return V. |
| Algorithm 2: Keep Ranked Volume |  |
|  | Input: V<sup>i</sup> ∈ {0,1}<sup>d×h×w</sup>, k ∈ Z |
|  | Output: V<sup>o</sup> ∈ {0,1}<sup>d×h×w</sup> |
| 1 | Find and count the connected components: V<sup>cc</sup> ← bwconncomp(V<sup>i</sup>, 26); /* ‘bwconncomp’ is a MATLAB function */ |
| 2 | Creates a label matrix corresponding to each connected component: L<sup>cc</sup> ← labelmatrix(V<sup>cc</sup>); /* ‘labelmatrix’ is a MATLAB function */ |
| 3 | Calculate parameters of each component: AreaProp ← regionprops(V<sup>cc</sup>, ′Area′); /* ‘regionprops’ is a MATLAB function */ |
|  | Sort the areas: (·, index<sub>sort</sub>) ← sort(AreaProp, ′descend′); |
| 4 | Copy the volume: V<sup>o</sup> ← V<sup>i</sup>; |
| 5 | If k > 0 do: |
| 6 | Remove the components other than the k<sup>th</sup> largest volume: V<sup>o</sup>[L<sup>cc</sup>! = k] ← 0; |
| 7 | Else do: |
| 8 | Remove the components of the k<sup>th</sup> largest volume: V<sup>o</sup>[L<sup>cc</sup> == k] ← 0; |
| 9 | End |
| 10 | Return V<sup>o</sup>. |
| 11 |  |
| Algorithm 3: Serial Dilate & Erode |  |
|  | Input: V<sup>i</sup> ∈ {0,1}<sup>d×h×w</sup>, [t<sub>1</sub> ⋯ t<sub>k</sub>] ∈ Z<sup>k</sup> |
|  | Output: V<sup>o</sup> ∈ {0,1}<sup>d×h×w</sup> |
| 1 | Set the parameter size of structure element: size; |
| 2 | Initialize a structure element: se ← strel(′cube′, size); /* ‘strel’ is a MATLAB */ |
| 3 | Initialize the volume: V<sup>o</sup> ← V<sup>i</sup>; |
| 4 | Foreach i ∈ [1, k] ∩ Z do: |
| 5 | Foreach j ∈ [1, abs(t<sub>i</sub>)] ∩ Z do: |
| 6 | If t<sub>i</sub> > 0 do: |
| 7 | Dilate the volume to fill the hole inside: V<sup>o</sup> ← imdilate(V<sup>o</sup>, se); |
| 8 | Elseif t<sub>i</sub> < 0 do: |
| 9 | Erode the volume to fit a specific size: V<sup>o</sup> ← imerode(V<sup>o</sup>, se); |
| 10 | End |
| 11 | End |
| 12 | Return V<sup>o</sup>. |
| Algorithm 4: Count Area & Ratio |  |
|  | Input: I<sup>i</sup> ∈ R<sup>h×w</sup>, τ ∈ R, n<sub>conn</sub> ∈ Z, M<sup>i</sup> ∈ {0,1}<sup>h×w</sup> |
|  | Output: M<sup>r</sup> ∈ {0,1}<sup>h×w</sup>, a<sup>r</sup> ∈ R<sup>m</sup>, r<sup>r</sup> ∈ R<sup>m</sup> |
| 1 | Normalize the input image: I<sup>o</sup> ← (I<sup>i</sup> − I<sup>i</sup>)/(I<sup>i</sup> − I<sup>i</sup> + 10<sup>−5</sup>); |
| 2 | Filter the image using a high passing filter: I<sup>o</sup> ← (1 − G) ∗ I<sup>i</sup>; /* G is a gaussian filter kernel function */ |
| 3 | Get the areas of the red channel: M<sup>r</sup> ← (I<sup>o</sup>[⋯,1] > τ[1])&(I<sup>o</sup>[⋯,2] < τ[2])&(I<sup>o</sup>[⋯,3] < τ[3]); |
| 4 | Find and count the connected components: M<sup>cc</sup> ← bwconncomp(M<sup>i</sup>, 8); /* ‘bwconncomp’ is a MATLAB function */ |
| 5 | Creates a label matrix corresponding to each connected component: L<sup>cc</sup> ← labelmatrix(M<sup>cc</sup>); /* ‘labelmatrix’ is a MATLAB function */ |
| 6 | Calculate parameters of each component: AreaProp ← regionprops(M<sup>cc</sup>, ′Area′); /* ‘regionprops’ is a MATLAB function */ |
| 7 | Get the number of components: m ← \|AreaProp\| − 1; |
| 8 | Initialize the areas and ratios for each component: a<sup>r</sup> ← [0]<sub>m×1</sub>, r<sup>r</sup> ← [0]<sub>m×1</sub> |
| 9 | Foreach i ∈ [1, m] ∩ Z do: |
| 10 | a<sup>r</sup>[i] ← ∑(L<sup>cc</sup> == i + 1) ∩ M<sup>r</sup>; |
| 11 | r<sup>r</sup>[i] ← a<sup>r</sup>[i]/∑(L<sup>cc</sup> == i + 1); |
| 12 | End |
| 13 | Return M<sup>r</sup>, a<sup>r</sup>, r<sup>r</sup>. |

